# ACC neural ensemble dynamics are structured by strategy prevalence

**DOI:** 10.1101/2022.11.17.516909

**Authors:** Mikhail Proskurin, Maxim Manakov, Alla Y. Karpova

## Abstract

Medial frontal cortical areas are thought to play a critical role in the brain’s ability to flexibly deploy strategies that are effective in complex settings. Still, the specific circuit computations that underpin this foundational aspect of intelligence remain unclear. Here, by examining neural ensemble activity in rats that sample different strategies in a self-guided search for latent task structure, we demonstrate a robust tracking of individual strategy prevalence in the anterior cingulate cortex (ACC), especially in an area homologous to primate area 32D. Prevalence encoding in the ACC is wide-scale, independent of reward delivery, and persists through a substantial ensemble reorganization that tags ACC representations with contextual content. Our findings argue that ACC ensemble dynamics is structured by a summary statistic of recent behavioral choices, raising the possibility that ACC plays a role in estimating – through statistical learning – which actions promote the occurrence of events in the environment.

## Introduction

Flexibility in choosing one’s behavioral strategy is a foundational characteristic of intelligent behavior, enabling rapid detection and adaptation to changes in the environment. In mammals, the brain’s ability to adaptively change behavior is thought to depend on the coordinated action of medial frontal cortical areas that keep track of the information necessary for context-appropriate choices of strategy (Domenech et al., 2020; Donoso et al., 2014). Functional imaging studies in human subjects and non-human primates have suggested that the anterior cingulate cortex (ACC), in particular, is a critical cortical node that translates contextual information into strategy changes (Domenech et al., 2020; Donoso et al., 2014; Hayden et al., 2011; Kolling et al., 2012; Sarafyazd and Jazayeri, 2019; Seo et al., 2014). Less clear is what specific computations are instantiated in the neural ensemble dynamics, in part because analyses of the incredibly diverse neural responses in ACC have yielded few simplifying principles. Several conceptual interpretations of the observed diversity of individual neural responses in frontal cortical regions such as the ACC have been put forth, including a direct role in enabling the separation of distinct contexts (Rigotti et al., 2010), and in tracking the animal’s evolving motor state (Musall et al., 2019; Stringer et al., 2019). However, novel yet interpretable changes in ACC responses associated with behavioral choices other than those explicitly instructed or prompted by experimental manipulations have also been recently observed in behavioral frameworks that relaxed some of the experimental control over the subjects’ behavioral responses (Schuck et al., 2015; White et al., 2019), raising the possibility that additional organizing principles for the ACC ensemble dynamics remain to be discovered.

One notable early observation related to the specificity of ACC’s functional engagement for settings with less external instruction highlighted a particularly marked enhancement of this cortical region’s hemodynamic signal during self-guided exploration, with both the self-generation of a specific information-sampling strategy and its evaluation contributing to the observed signal (Walton et al., 2004). The advantage of choosing one’s strategy for sampling information when learning in complex settings has long been appreciated(Markant and Gureckis, 2014); indeed, the idea that subjects learn more effectively when they self-direct their learning experience is even central to many educational philosophies (Boekaerts, 1997; Bruner, 1961). Implicit in these notions is a key role for the self-guided component of knowledge acquisition. As such, it is notable that the medial frontal lobe, including the supplementary motor cortex (SMC) and the ACC, have been proposed to process information related to self, and to implement flexible, self-guided actions (reviewed in (Passingham et al., 2010), but see (Schüür and Haggard, 2011)). The cingulate region has also begun to feature prominently in the emerging elegant efforts to experimentally expose task-related information seeking and to uncover its neural underpinnings (Wang and Hayden, 2020; White et al., 2019). Thus, evaluating neural activity in behavioral settings that do not prompt specific responses or provide explicit instruction regarding task-related strategies may represent a fruitful direction in the search for potential organizing principles of the frontal cortical neural ensemble dynamics.

In this study, we evaluate medial frontal cortical ensemble activity in the context of a behavioral framework that requires rats to search, without any explicit instruction, for latent structure within a space of action sequences to uncover the preferentially rewarded one. Taking advantage of the observation that under such task conditions, rats continue to periodically sample alternative sequences even having discovered the latent target, we demonstrate that in the ACC, ensemble activity associated with a specific action sequence is markedly re-organized between inferred global task contexts, yet invariably tracks the local prevalence of that sequence in the animal’s behavioral choices. Prevalence encoding in the ACC is robust, accounting for a substantial fraction of activity variance during sequence execution in over half of ACC neurons. Although few individual ACC neurons display modulation that spans the entire temporal extent of sequence execution, information about the prevalence in recent behavioral history of the currently executed sequence is maintained at the ensemble level. Strikingly, prevalence encoding is featured particularly strongly in the underexplored, most rostral portion of the ACC with homology to primate area 32D. Our findings demonstrate that the ACC network enables a continuous representation of strategy prevalence through a substantial degree of reorganization that tags these representations with global contextual content, and suggest that long-term tracking of strategy prevalence might be a fundamental part of the animals’ algorithmic approach to complex settings.

## Results

### Behavioral framework for self-guided search for task structure and strategy encoding in ACC

To examine frontal cortical neural dynamics during self-guided exploration of a complex environment, we developed a behavioral framework that requires rats to search, without any explicit instruction, a structured space of action sequences to infer a specific latent pattern that is preferentially rewarded. A typical behavioral session in this framework comprises a series (750-2000) of self-initiated trials that involve a choice between two options – left or right nose port – with a reward being delivered upon execution of the latent (i.e. not explicitly revealed) target sequence of choices – such as right-right-left, ‘RRL’. Our behavioral paradigm requires center-port entries between each choice of a side port. Hence, execution of the length 3 sequence ‘RRL’ requires the series of 6 port entries: cRcRcL (Fig. 1a). The requirement for center port entries between each side port entry was chosen in part to control the movements associated with the execution of a given sequence; with this requirement, the rat’s movements to select each instance of ‘R’ in the sequences ‘RRL’, ‘LLR’, etc., are behaviorally constrained to involve withdrawing from the center port, then moving to and entering the right port. This center port requirement, therefore, facilitates the separation of neural signals specifically associated with the sequence of ‘R’ and ‘L’ choices from the movements associated with selecting them.

**Fig. 1.**
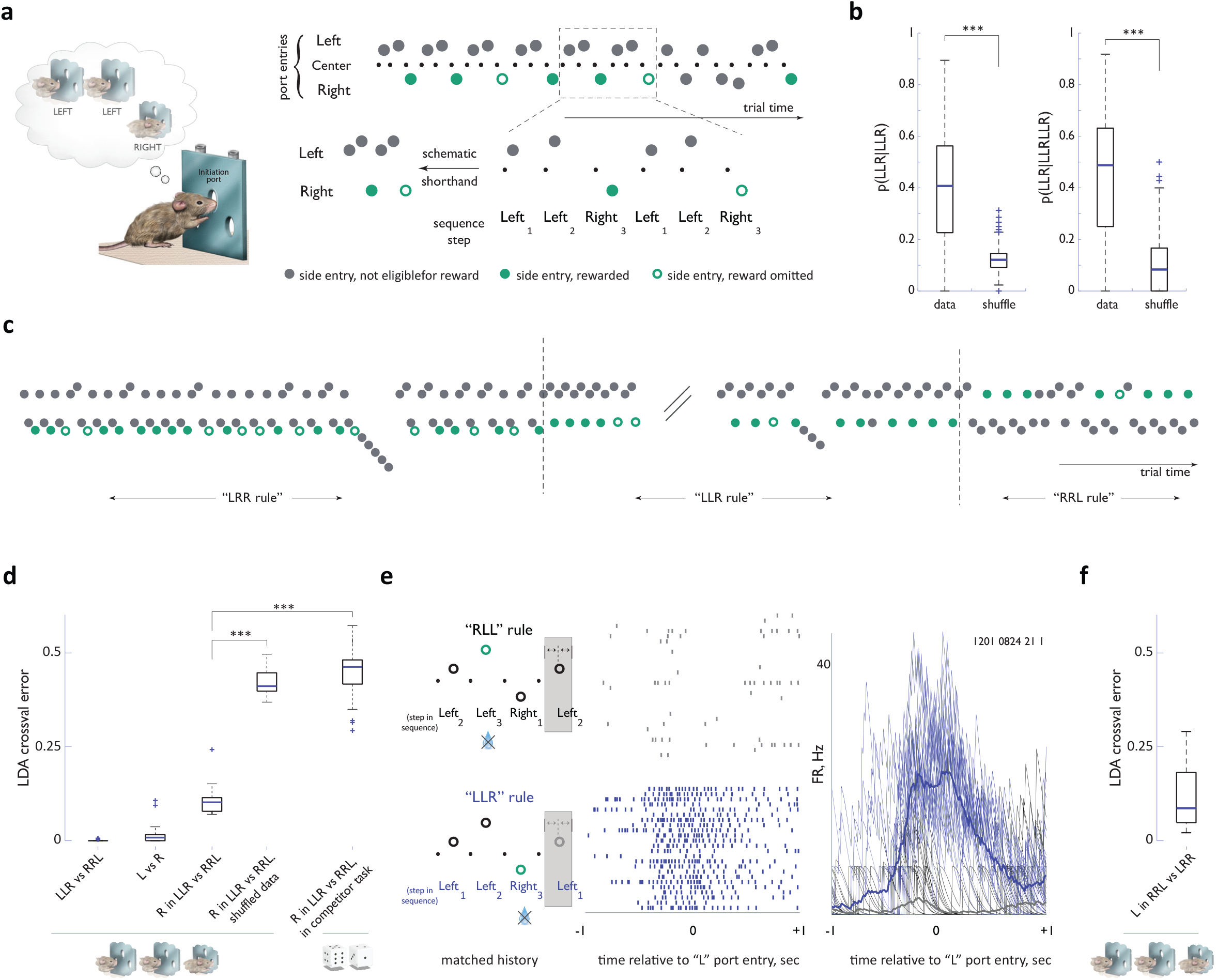
Strategy encoding in the ACC. **a. Left panel:** Concept of the behavioral task. After initiating at the center port, the animal is eligible to receive a reward only if his sequence of past choices conforms to a latent target sequence, like ‘Left-Left-Right’. Note that the identity of the latent target is not otherwise cued in any way. **Right panel:** Schematic of the notation used for behavioral data presentation. Note that nose port entries are omitted from schematics in other panels for simplification. **b**. Probability of target sequence concatenations across the behavioral dataset. **c**. Sample behavioral trace, in trial time, around two block transitions. **d**. Cross-validated performance errors for linear classifiers trained to distinguish ACC representations of different strategy components. Rolling dice indicate virtual competitor sessions(Barraclough et al., 2004; Tervo et al., 2014). n=36 sessions, N= 4 animals for all sequence task comparisons; n=9 sessions, N=3 animals for competitor sessions. **e**. Comparison of activity traces for an example ACC neuron in the 1-second window around entry into the left nose port within the ‘LLR**L**’ context also matched in reward history during either ‘RLL’ rule (black) or ‘LLR’ rule (blue). **f**. Cross-validated performance error for a linear classifier trained to distinguish ACC representations of ‘R’ in ‘LLR’ vs ‘RLL’ contexts. n=11 sessions, N=1 animal. ***, p < 0.001.

To ensure that our animals continue to self-direct a search for the relevant task structure throughout each behavioral session, we changed the latent target sequence identity in an unsignalled manner every 250-500 trials (Methods). To collect a sizable amount of reward in this task, an animal needs to locally structure its choices to preferentially conform to the target sequence, yet remain flexible enough to efficiently adapt to unsignalled changes in its identity (Fig. 1a,b). For most experiments in this study, we restricted the set of possible latent target patterns in any individual session to some, or all, of the four non-trivial three-step sequences (‘LLR’, ‘RRL’, ‘LRR’ and ‘RLL’). Under these conditions, expert animals flexibly and reliably discovered and exploited the locally-relevant latent target: within tens of trials following block transitions, the animals’ choices typically became dominated by patterns conforming to the new target sequence even in the presence of substantial (up to 30%) sporadic omission of reward for the properly executed target sequence (Figs. 1b-c, 2b, Supplementary Fig. 1). This marked behavioral flexibility did not result from an inability to commit to the discovered target sequence. Indeed, the target sequence clearly dominated behavioral choice streams during blocks of stability, with long concatenations of the target sequence frequently evident in the behavioral records (Figs. 1b-c,2b, Supplementary Fig. 2). Nevertheless, clear deviations from this dominant pattern, with the animals’ choices appearing instead to conform to other possible target sequences, were also present within all blocks (Figs. 1c, 2a-c). Such transient deviations away from the dominant sequence were present even when no reward was omitted for its correct execution (Supplementary Fig. 2), suggesting that animals naturally continue to sporadically sample other sequences even when not obviously extrinsically prompted. Thus, with only two well-defined individual actions (‘left’ and ‘right’), this framework provides a means to engineer a rich and flexible repertoire of multi-step sequences that animals evaluate in various behavioral contexts.

ACC is thought to play a central role in motivating extended, multi-step behaviors (Holroyd and Yeung, 2012), and our previous work demonstrated the rodent ACC homologue’s involvement in unguided discovery of action sequences (Tervo et al., 2021). Nevertheless, we first verified that the specific choice of strategy in this more complex sequence task is reflected in ACC neural dynamics by establishing that individual sequences could be decoded from ensemble activity. We initially focused on contrasting the encoding of ‘LLR’ and ‘RRL’ – the two main target sequences in our dataset. A simple classifier performed with near perfect accuracy on the task of distinguishing the two representations (Figure 1d, median cross-validated classification error 0.0, IQR 0.006, Methods). However, any interpretation of the decodability of the full ‘LLR’ vs ‘RRL’ ACC representations is confounded by a strong difference in the encoding of the individual steps composing the sequences (‘L’ vs ‘R’) (Figure 1d, median cross-validated classification error 0.008, IQR 0.016). This prompting us to examine more selectively whether the encoding of the same action, such as ‘R’, differs between the putative ‘LLR’ and ‘RRL’ sequences. To control for a possible influence of the local choice context unrelated to deliberate sequencing, we performed an equivalent analysis for all ‘LLR’s and ‘RRL’s found in the choice streams generated in a competitive setting that pushes animals to abandon strategic stringing together of individual choices into action sequences (Barraclough et al., 2004; Tervo et al., 2014). The classification error for this comparison was significantly lower when animals had to preferentially follow specific sequential strategies than when trained to avoid sequential patterns in their behavior (Fig. 1d, median cross-validated error 0.10, IQR 0.038 for the sequence task vs mdian error 0.41 and IQR 0.05 for the competitive setting), suggesting that at least in part, the differential encoding of ‘R’ in ‘LLR’ vs ‘RRL’ reflects its immersion in distinct strategies. Finally, we observed a similarly marked difference in the encoding of ‘L’ between circularly permuted strategies (‘RRL’ and ‘LRR’), even when reward was omitted for correct sequence execution thus ensuring full matching of immediately preceding history of choice/reward pairings (Figs. 1e,f). Combined, these observations argue that individual multi-step sequential strategies have distinct ACC representations and suggest that evaluation of broader contextual modulation should be restricted for different instances of the same sequence execution.

### Identification of distinct sequence instances in the behavioral stream

The robust performance our rats displayed on this task is consistent with the notion that settings requiring subjects to discover latent structure through self-guided exploration align particularly well with the way brains naturally make sense of complex environments (Gottlieb and Oudeyer, 2018; Tervo et al., 2016; Wang and Hayden, 2021). However, such an experimental framework poses challenges for parsing the behavioral record to identify ‘legitimate’ sequence instances. Parsing a continuous stream of left and right choices to identify ‘legitimate’ sequences is easy for the ‘dominant’ condition where a high prevalence of sequence concatenation, and a scarcity of choices that conform to other patterns, argue that almost every instance of a pattern conforming to the target sequence is likely to be one actually evaluated by the animal (see Methods for a detailed description of the filter used to select ‘dominant’ sequence instances). In contrast, parsing is much harder for the ‘exploratory’ condition outside of clear sequence concatenations. For example, although some ‘RRL’ patterns within runs of ‘LLRRL’ in a ‘LLR’ block might reflect a true pairing of the locally dominant pursuit of ‘LLR’ with a quick exploratory evaluation of ‘RRL’ tagged on in a manner akin to strategy mixing (Donoso et al., 2014), most of such instances likely reflect a mere apposition of the ‘LLR’ and the ‘RL’ sequences. We therefore next sought to delineate an objective criterion for including any lone putative exploratory sequence instance in the subsequent analyses, focusing on sessions that contained mostly ‘LLR’ and ‘RRL’ blocks.

Our approach was grounded in the expectation that animals would pause, if only briefly, at the side nose port on the last step of a true exploratory sequence instance, marking sequence completion. Under this assumption, putative exploratory sequence instances for which the duration of the third step exceeds a preset threshold, can be objectively included in the ‘exploratory’ dataset. The presence of clear shifts to longer within-side-port and side-to-side durations for step three – as compared to steps one and two – across all ‘dominant’ sequence instances for which the otherwise scheduled reward was omitted (Supplementary Fig. 3, Methods) permitted an unbiased selection of the specific threshold. The ‘exploratory’ dataset thus included sequence concatenations and lone sequence instances that passed the temporal selection threshold (see Methods for the full description of the selection procedure). Overall, the rich contextual variation associated with the execution of different instances of any specific target sequence in both ‘dominant’ and ‘exploratory’ conditions provided us with an opportunity to determine whether examining frontal cortical neural activity through the lens of this natural contextual variation might reveal dynamics that would shed additional light on the computations performed by these circuits.

### Functional reorganization of the ACC network constrains its representation of a specific strategy in distinct inferred global task contexts

ACC is thought to guide contextually-appropriate strategy selection (reviewed in (Heilbronner and Hayden, 2016; Holroyd and Verguts, 2021; Kolling et al., 2016; Monosov et al., 2020; Shenhav et al., 2016)), prompting us to begin our analysis of task-related neural responses by evaluating how global behavioral context defined by which sequence has come to locally dominate the animal’s choices – likely a manifestation of the animal’s inference about the currently relevant task structure – is reflected in ACC ensemble dynamics. Specifically, we first sought to determine if ACC activity associated with a specific behavioral sequence is reorganized depending on whether that sequence represents a dominant behavioral strategy or is sampled as a part of an exploratory bout. Indeed, while abrupt changes in the activity of ACC neurons coincident with behavioral transitions to exploration have been reported previously (Durstewitz et al., 2010; Emberly and Jeremy, 2019; Karlsson et al., 2012; Powell and Redish, 2016), whether these transitions in ACC activity merely mark the behavioral state change, or reflect an actual task-related re-organization of activity whereby representations of individual strategies are also marked with contextual content, remains unclear. Targeted recordings of ACC activity – performed in a wireless configuration that did not impair the animals’ behavioral flexibility – revealed that marked differences in activity associated with the execution of specific sequences in ‘dominant’ versus ‘exploratory’ contexts could indeed be readily observed across many ACC neurons, frequently evident even in non-trial averaged activity traces (Fig. 2d). To facilitate detailed comparisons of neural activity in the face of inevitable variability in the spatio-temporal profiles of movements between nose ports for different instances of sequence execution, we have focused all analyses on five 500 millisecond windows anchored on center and side noseport entry events associated with the individual steps in the sequence (Fig. 2a, see also a more detailed evaluation of potential motor confounds below).

**Fig. 2.**
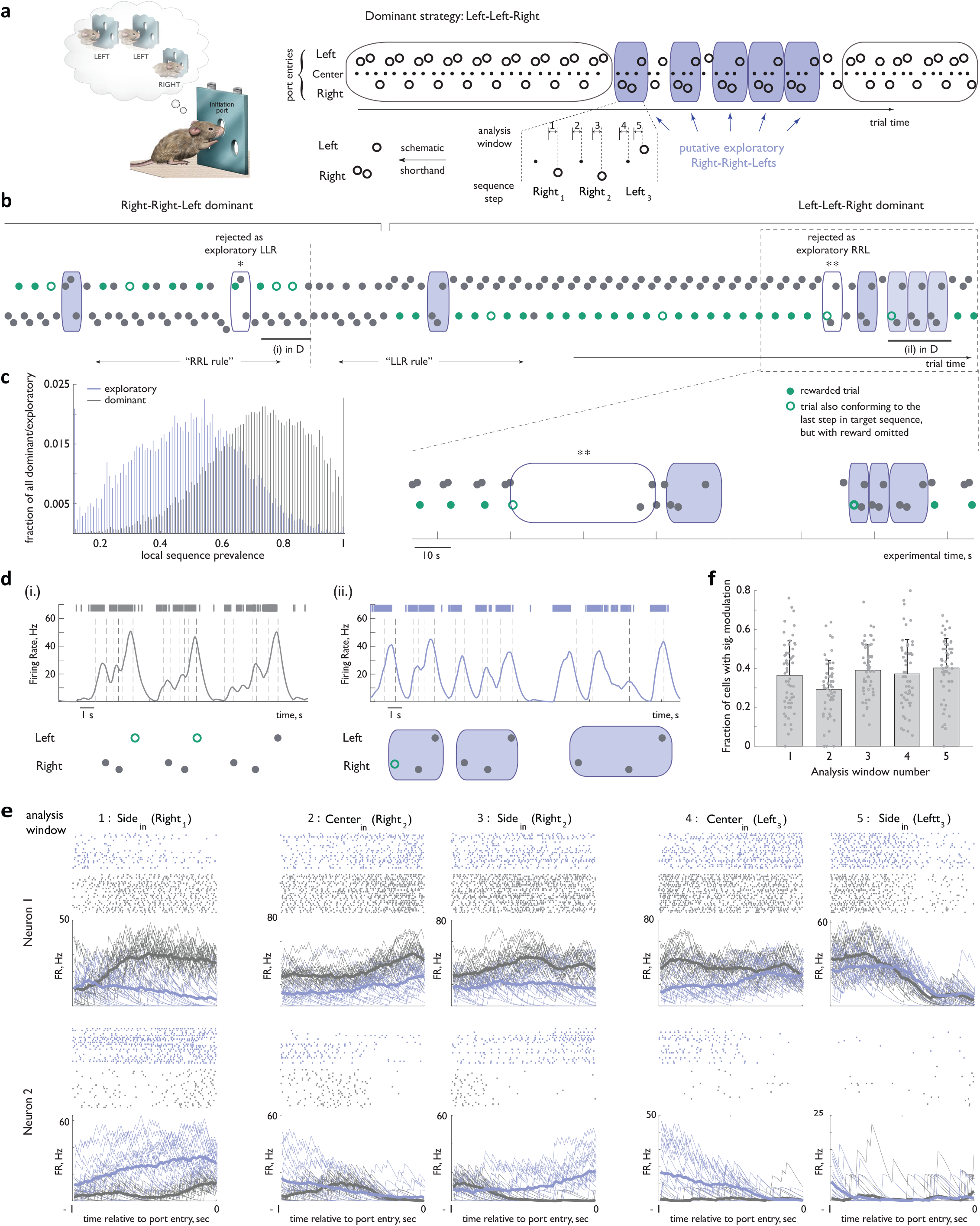
Activity of ACC neurons associated with a specific sequence of actions changes depending on whether that sequence represents the dominant strategy or is a transiently re-explored alternative. **a. Left panel:** Concept of the behavioral task (same as in Figure 1). **Right panel:** (Top) Block-wise structure promoted the local pursuit of one dominant sequence (here, ‘Left-Left-Right’, as shown in the thought bubble) at the expense of others. The dominant strategy was occasionally interrupted by explorations of alternative sequences (here, likely ‘Right-Right-Left’, blue shading). (Bottom) Five analysis windows chosen to minimize trajectory confounds were anchored on center- and side- noseport entry events associated with each of the three steps in the sequence, with the exception of the first center port entry. The latter omission was chosen to minimize the contribution from feedback-related activity modulation associated with preceding choices. **b. Top panel:** Sample behavioral trace, in trial time, around a block transition. **Bottom panel:** boxed region of the behavioral trace, in trial time. Note a marked preference at the beginning of the behavioral trace for ‘Right-Right-Left’ and at the end for ‘Left-Left-Right’. Putative exploratory sequences are shaded blue (see Methods for details about how exploratory sequences were identified). *, rejected as an exploratory sequence because it overlapped with the dominant sequence to its left, and did not have the temporal profile consistent with sequence marking (see text) to be rescued from the ‘discard’ group; **, rejected as a putative exploratory sequence because of an overlap with the dominant sequence to its left, and an unusually long break between putative steps 1 and 2. **c**, Distribution of local sequence prevalence values for ‘Left-Left-Right’ and ‘Right-Right-Left’ across all dominant and exploratory instances in implanted animals. **d**. Activity of an example ACC neuron for three concatenated unrewarded instances of ‘Right-Right-Left’ at the end of the dominant epoch in **(b)** (**left panel**) and during an exploratory bout following subsequent dominance of ‘Left-Left-Right’ (**right panel**). **e**. Activity of two other ACC neurons from the behavioral session in **(b)**, aligned in the five one-second analysis windows anchored on port entry events. Grey: dominant instances of ‘Right-Right-Left’ that followed an unrewarded ‘Right-Right-Left’. Blue: exploratory instances of ‘Right-Right-Left’. **f**. Fraction of all recorded ACC units that displayed a significant modulation between ‘dominant’ and ‘exploratory’ contexts for each of the five analysis windows. Individual points correspond to different sessions. Error bars represent standard deviation. n=35 sessions, N=4 animals.

An alignment of sequence-related activity in the five constrained analysis windows across different sequence instances exposed a variety of contextual modulation of activity in individual cells (see Fig. 2e for alignment of spike rasters, Fig. 3b for alignment of heatmap representations). Indeed, both decreases, increases and mixed modulations of activity in the exploratory context relative to that observed when the sequence represented the dominant strategy were observed (see Figs. 2e, 3a for examples). At least 76 % of all recorded units (787 of 1042 total ACC recorded units, with no separation of potential interneurons) displayed significant modulation in at least one of the five analysis windows – a lower bound given the limitation that the modest size of the exploratory dataset places on the statistical power of these analyses – and a roughly equal fraction of all units displayed modulation in each of the analysis windows (Fig. 2f). Thus, activity changes related to the inferred global task context are present at the single-cell level in many ACC neurons.

**Fig. 3.**
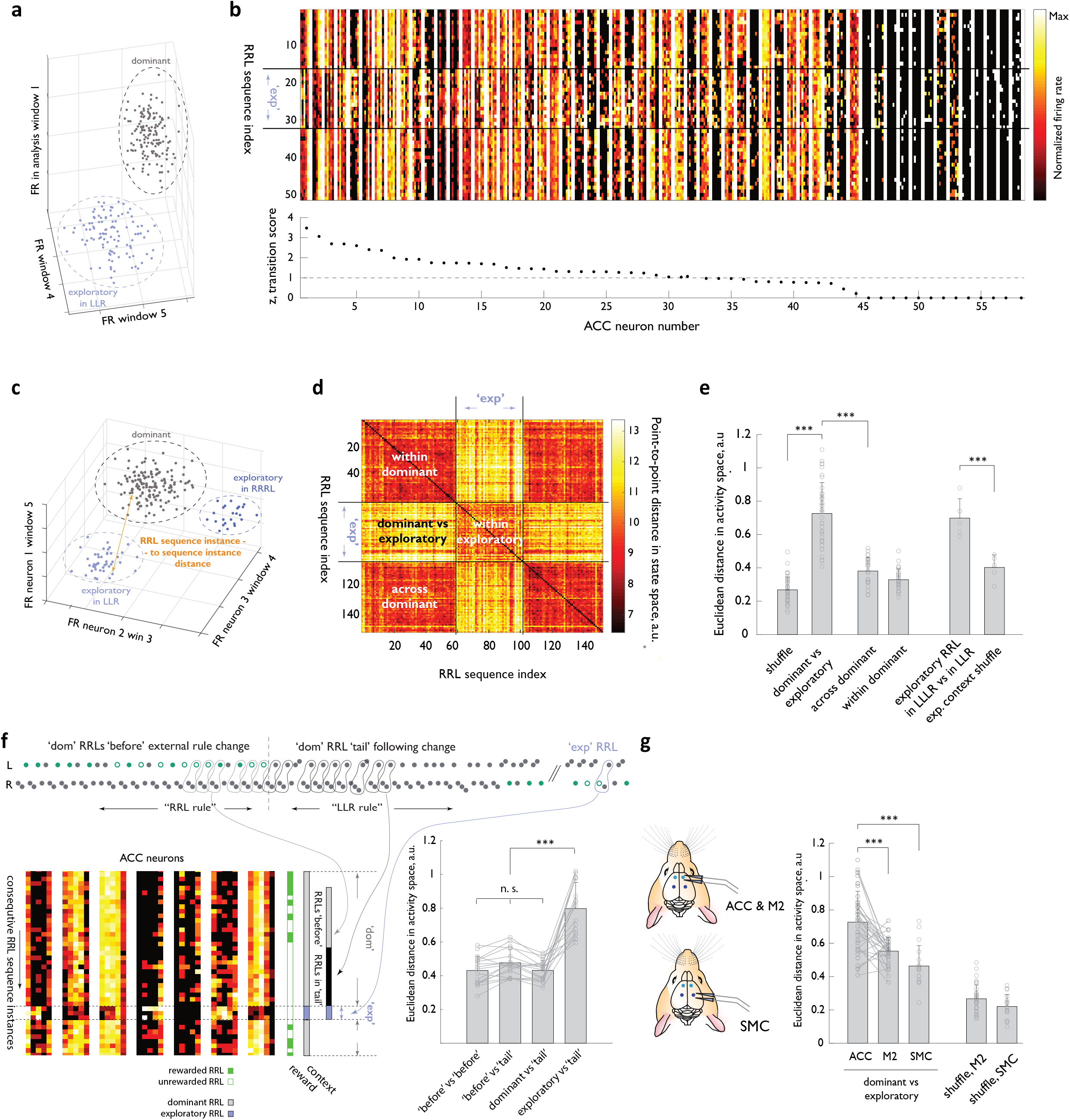
Representational transitions in ACC reflect large-scale functional reorganizations of the ACC network between inferred global behavioral contexts. **a**. Schematic of the activity state space for an individual neuron. Three of the five dimensions corresponding to the analysis windows are shown. Two clouds schematize activity of that neuron associated with a specific behavioral sequence for all dominant (grey) and exploratory (blue) instances of the sequence. **b**. Heat map representations of normalized activity associated with ‘Right-Right-Left’ sequence execution for 58 simultaneously recorded ACC neurons. Different sequence instances are stacked vertically, with two ‘dominant’ blocks separated by a period when the ‘Right-Right-Left’ sequence was occasionally explored in the background of ‘Left-Left-Right’ dominance. ‘exp’, ‘exploratory’ instances. Neurons are arranged according to a ‘transition score’ defined as the distance between the two cloud centroids normalized by root mean of variance within each cloud (see Methods). **c**. Schematic of the ensemble state space. Each of the simultaneously recorded neurons contributes five dimensions. Only three dimensions are shown for simplicity. Two distinct clouds show ‘exploratory’ instances of RRL in the context of distinct ‘dominant’ sequences, LLR and LLLR. **d**. Similarity matrix for an example session, comparing the ACC ensemble activity across ‘Right-Right-Left’ instance pairs, using Euclidean distance in the network state space. Black lines indicate the boundary between ‘dominant’ and ‘exploratory’ contexts. **e**. Euclidean distance (RRL instance-to-instance) in the state space between the relevant ‘dominant’ and ‘exploratory’ clouds for the experimental ACC data, and for the control state space, where the labels of ‘dominant’ and ‘exploratory’ (or of the specific ‘exploratory’ context) were randomly shuffled across the dataset. n=35 sessions, N=4 ACC-implanted animals. **f**. Behavior of the ACC ensemble during persistence of the dominant strategy past the unsignalled transition in the rewarded target. **Top panel:** example behavioral transition with a long dominant RRL ‘tail’. **Bottom left panel:** heat-map representation of the activity of 5 ACC cells for consecutive RRL instances before (i.e. when RRL dominance coincided with it being the rewarded target, ‘before’ in panel), during (when RRL continued to dominate the animal’s choices but the target sequence had changed, ‘tail’ in panel) and after the behavioral transition. Note that the latter set of examples included several exploratory instances (executed much later, once another sequence had established dominance, ‘exp’ in the panel), as well as several of the subsequent instances of ‘dominant’ RRL later in the session. ‘exp’, ‘exploratory; ‘dom’, ‘dominant’. **Bottom right panel:** RRL instance-to-instance Euclidean distance in the state space across these distinct epochs for all long ‘dominant tail’ examples. n=12 sessions, N=4 animals. **g**. Euclidean distance between the centroids of the ‘dominant’ and ‘exploratory’ clouds for the experimental ACC, M2 and SMC data, and for the control state spaces, where the labels of ‘dominant’ and ‘exploratory’ were randomly shuffled across the dataset. n=37 sessions, N=4 ACC/M2 -implanted animals; n=18 sessions, N=3 SMC-implanted animals. Error bars represent standard deviation. n.s., not significant, ***, p < 0.001.

To examine these contextual changes in the ACC representation associated with a specific behavioral sequence in more detail, we next evaluated population responses in a neural state space. For these analyses of potential contextual re-configurations, we used a state-space framework that assigns the firing rate of an individual neuron in each of the five analysis windows to a single dimension. Consequently, a point in this five-dimensional state captures the activity of that neuron during one instance of sequence execution (Fig. 3a-b, Methods). The ensemble representation, in turn, is captured by further expanding the state space dimensionality to include these five dimensions for each of the simultaneously recorded neurons (Fig. 3c). When we used similarity matrices to visualize instance-to-instance distance between ACC representations for a specific sequence in the full state space, a substantial separation of the two clusters formed by the ‘dominant’ and ‘exploratory’ instances of sequence execution was readily evident (Fig. 3d). Indeed, the Euclidean distance between the centroids of these experimentally observed ‘dominant’ and ‘exploratory’ groups differed markedly from that observed when the ‘dominant’ and ‘exploratory’ labels were randomly shuffled across the set of all instances of sequence execution (Fig. 3e, 0.73 ± 0.18 for experimental data vs 0.268 ± 0.077 for shuffled data, p< 10^−12^, Wilcoxon rank-sum test, n=35 sessions, 4 animals). Moreover, the mean Euclidean distance in the activity state space between two ‘dominant’ blocks separated in time was significantly smaller than the mean pairwise distances between a dominant and exploratory block, arguing against the possibility that the observed representational transitions in ACC arose from an instability in neural recordings (Fig. 3 d,e). Thus, ACC ensemble activity markedly changes its representation of a particular behavioral sequence when that sequence no longer represents the locally dominant behavioral strategy but settles back into a similar state every time the animal returns to its pursuit of that sequence over all others.

The strong similarity of ACC ensemble configurations across distinct, temporally segregated ‘dominant’ contexts (Fig. 3e) argues that these representational rearrangements reflect something other than a mere episodic record of distinct behavioral contexts. Given that ACC is thought to encode signals that convey the value of pursuing alternative courses of action as opposed to the current, default action plan (Behrens et al., 2007; Blanchard and Hayden, 2014; Hayden et al., 2009; Karlsson et al., 2012; Kolling et al., 2012; Kolling et al., 2014; Ma et al., 2016; McGuire et al., 2014; O’Reilly et al., 2013; Powell and Redish, 2016; Procyk et al., 2000; Schuck et al., 2015; Tervo et al., 2021), the abrupt transitions in the ACC representation of specific sequences we observed may be designed to tag these representations simply as ‘dominant’/’default’ or ‘exploratory’/ ‘alternative’. However, broader contextual signals are also thought to be present in this cortical region (Caracheo et al., 2018; Euston et al., 2012; Seamans and Floresco, 2021; Shenhav et al., 2016; Tomlin et al., 2006), and thus the ACC representational transitions may carry a richer contextual content. To distinguish these possibilities, we compared, in a small subset of sessions, the ACC representation of exploratory ‘RRL’s in two separate contexts: the ‘LLR’ context (i.e., one where the animal otherwise pursued ‘LLR’ as the default strategy) and the ‘LLLR’ context. Despite the limited statistical power of exploratory datasets, representational changes between these distinct exploratory contexts were clearly evident both at the single neuron and the population level (Fig. 3e, Euclidean distance of 0.70 +/-0.11 for two sets of exploratory RRL representations vs 0.40 +/-0.08 for a control set with scrambled contextual labels, p< 0.008, Wilcoxon rank-sum test, n=5 sessions, 2 animals). Notably, the marked restructuring of ACC network configuration that underpins the differential encoding of exploratory ‘RRL’s in these two contexts argues against the possibility that these large scale-reconfigurations merely reflect the absence or presence of reward for a given sequence in that task epoch since none of the ‘RRL’s in *either* context were ever rewarded. Rather, these observations support the notion that the ACC network functionally reconfigures in distinct behavioral contexts and argues that the inferred global contextual content of representational transitions in ACC is richer than the ‘dominant’/’exploratory’ dichotomy.

Our parsing of the global behavioral context based on the dominant strategy chosen by the animals rather than based on the experimentally imposed identity of the target sequence is in keeping with the view that the medial prefrontal cortex does not simply track external cues that situate the animal in time and place, but instead reflects the state of the animal’s emotions and beliefs (Caracheo et al., 2018; Euston et al., 2012; Seamans and Floresco, 2021). Nevertheless, the externally imposed task context (i.e., the identity of the latent target sequence at any moment) and its parsing by the animal that presumably shapes the choice of the dominant strategy are strongly correlated in expert animals. Thus, to further probe the validity of parsing based on the animal’s behavior, we next examined more closely the ACC neural ensemble activity evolution in cases where the dominance of a specific sequence persisted in the face of a change in the identity of the rewarded target (Fig. 3f). Ensemble states associated with sequence execution during such dominant sequence ‘tails’ did not cluster away from ones observed earlier in the ‘dominant’ context, and were equally distant from the exploratory cluster (Fig 3f, Euclidean distance of 0.43 +/-0.06 for sequence instances within ‘tails’ to other dominant sequence instances vs 0.80 +/-0.15 for sequence instances within ‘tails’ to exploratory sequence instances, p< 10^−6^, Wilcoxon rank-sum test, n=17 ‘tail’ examples, N=12 sessions). Together with our previous observation that the ACC neural representation associated with a specific probabilistically rewarded action differed markedly depending on the strategy-related context (Tervo et al., 2021), the unambiguous co-segregation of the ensemble representation for sequence instances executed during even the longest post-target-sequence-switch ‘tails’ we observed here further argues against the simple ‘reward’/’no reward’ explanation of network reconfigurations. More broadly, these observations support parsing of the behavioral context through the lens of the selected strategy.

We also investigated whether the contextual reorganization of ensemble activity we observed in the ACC reflects an ACC-specific computation or is present in other parts of the frontal cortex. Indeed, neural correlates of decision task parameters have long been observed across many frontal cortical areas (Cisek, 2012; Cisek and Kalaska, 2010; Hunt and Hayden, 2017; Yoo and Hayden, 2018), and recent advances in the ability to simultaneously record from tens of thousands of neurons across many brain areas in the mouse have only further emphasized how widely distributed the task-related encoding can be in the brain (Steinmetz et al. 2018; Stringer et al. 2019). We focused on two distinct regions along the rostro-caudal axis of the medial lobe. At the rostral end, we targeted, within the same set of recording sessions, the immediately dorsal to the ACC part of the premotor cortex (henceforth, M2) that contains, among other areas, the putative rat homologue of the frontal orienting fields recently implicated in a range of sensory-guided decisions (Ebbesen et al., 2018). At the caudal end, we probed, in a separate set of animals, the neural dynamics in a premotor region that shares key afferent/efferent projection patterns with the primate SMC and is functionally required for self-guided action sequencing in our behavioral framework (Supplementary Fig. 4, Methods and (Manakov, Proskurin *et al, in preparation*); henceforth, SMC). Analysis of the M2 and the SMC ensemble representations associated with individual behavioral sequences over the course of long behavioral sessions revealed that although representational changes could also be observed in those parts of the medial lobe, these changes were significantly less pronounced and affected a smaller fraction of the recorded units, especially in the more caudal SMC (Fig. 3g, Supplementary Fig. 5). While we cannot rule out the possibility that the inherent sampling bias of the extracellular recordings has led us to miss some less active M2 or SMC neurons that also display a substantial degree of activity reorganization, these observations suggested that large-scale functional network re-arrangements between the inferred global behavioral contexts are particularly prominent in the ACC.

### Within each global functional re-configuration of ACC neural ensemble, the representation of a behavioral strategy is further shaped by its local prevalence

We next investigated whether ACC neural dynamics are further shaped by more local adjustments to the animal’s course of action. Indeed, within the otherwise largely stable functional ensemble configurations reflected in darker-colored squares of the similarity matrices (Fig. 3d, ‘within-dominant’ and ‘within-exploratory’ squares), some local variation in pixel intensity was consistently observed. Therefore, we sought to determine whether the underlying variability in the ACC neural ensemble dynamics characterizing individual instances of sequence execution may reflect local fluctuations in the animal’s choices. As a natural extension of defining global context (above) based on the roughly block-wise, persistent dominance of a single sequence, we chose the simplest summary statistic – sequence prevalence in recent choices – to capture local fluctuations in how much pursuit of the sequence in question is balanced by exploration of other strategies (Methods).

Firing rates indeed tracked local sequence prevalence in a substantial fraction of individual neurons that otherwise displayed a variety of response profiles (see Fig. 4a for examples). Although a fraction of ACC neurons tracked local sequence prevalence throughout sequence execution, the majority did so in a subset of the 5 analysis windows that were aligned to nose port entries, roughly equally distributed across the temporal extent of the multi-step sequence (Supplementary Fig. 6). To examine this modulatory influence on ACC activity in greater detail, we fit the relationship between the firing rates of individual neurons and the local sequence prevalence with a linear model – with or without the global context (‘dominant’/’exploratory’) as a fixed parameter – and evaluated the model’s performance through cross-validation (Methods). The model’s explanatory power was robust to the precise number of trials in the past used to calculate local sequence prevalence but was diminished if the estimate was shifted to largely comprise future trials (Supplementary Fig. 7). The cross-validated performance of the mixed-effects linear model that included global context exceeded the performance of the model that had prevalence as a single parameter (Supplementary Fig. 8), arguing that prevalence tracking and the large-scale functional reorganization of the ACC network between the ‘dominant’ and ‘exploratory’ contexts in our task reflect distinct computations. Strikingly, when we allowed the slope of the linear relationship between the strategy prevalence and the firing rate of individual neurons to vary depending on the global behavioral context, this simple behavioral summary statistic explained up to 80% of neural activity variance in individual ACC neurons during sequence execution (Fig. 4b,c; median 28, IQR 42 % of activity variance explained by prevalence across all recorded cells in the rostral ACC). The marked modulation of individual cell activity in the ACC by local strategy prevalence stands in contrast with a more modest modulation detectable in other parts of the medial lobe, in particular the SMC (Fig. 4b, Supplementary Fig. 8), pointing to a distinct role the rostral medial frontal cortical region plays in strategy contextualization. Within the ACC, a encoding potential gradient emerged following a detailed spatial reconstruction of the recorded units, with the most substantial modulation observed in the rostral-most portion thought to be homologous to the primate area 32D (Fig. 4b,c). Thus, this underexplored, rostral-most portion of the ACC may play a particularly prominent role in keeping a statistical representation of recent behavioral choices.

**Fig. 4.**
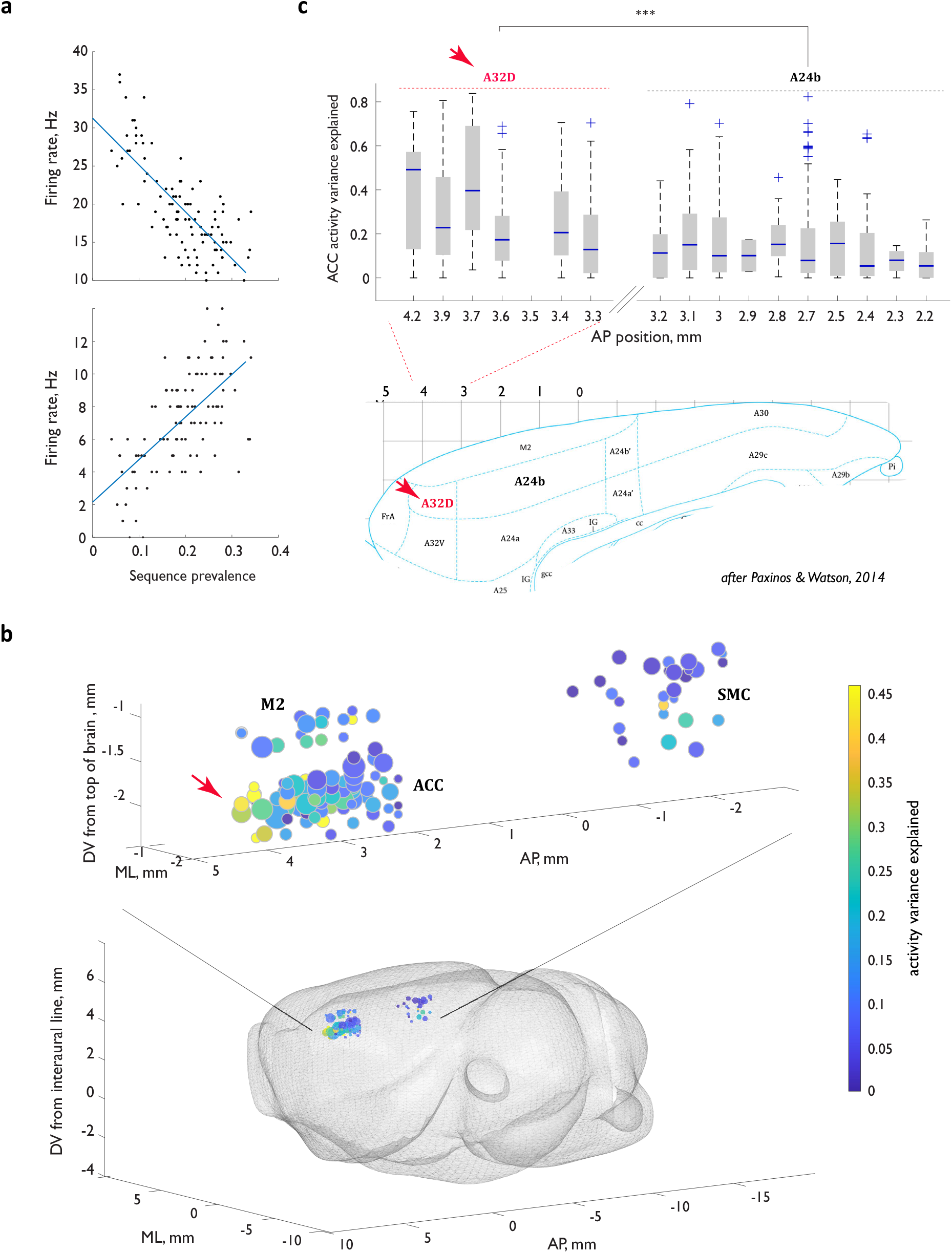
Activity of ACC neurons during sequence execution is markedly shaped by the local prevalence of the executed sequence. **a**. Firing rates as a function of local sequence prevalence for two example ACC neurons. **b**. Graphic representation of the explanatory power of the linear model across the spatial extent of the recording locations. **c**. Top panel: explanatory power in the rostral portion of the cingulate as a function of location along the anterior-posterior axis. Bottom panel: refence rat brain atlas section. Red arrow in **(b**,**c)** points to a cluster of the particularly strong model performance in the rostral portion of the cingulate that maps to the region homologous to the primate area 32D. For all box-and-whisker plots, central red line indicates the median, the bottom and top edges of the box indicate the 25th and 75th percentiles, respectively, and the whiskers extend to the most extreme data points not considered outliers. n.s., not significant; **, p<0.01; ***, p < 0.001.

Models of decision-making rarely include a summary statistic of the agent’s recent behavioral choices – such as the local strategy prevalence that we examine here. This prompted us to next consider whether the observed modulation of ACC ensemble activity by local strategy prevalence could be as easily explained by other, more commonly considered behavioral parameters. Indeed, if strategy prevalence at least to some extent reflects the animal’s local resolve to pursue that strategy, the detailed kinematics of the corresponding movement, the vigor of movement execution, or the associated reward expectation might plausibly co-vary with strategy prevalence. Furthermore, variation in strategy prevalence is necessarily accompanied by variation in detailed reward and choice history: known modulators of ACC activity. To investigate this issue, we thus asked how well linear models that either incorporate the more commonly considered parameters directly as regressors, or account for those parameters in an indirect way performed at explaining variance in ACC firing rates (see Methods).

Trial-by-trial measures of movement vigor and kinematics were directly quantified for each instance of sequence execution using sequence execution time and the first principal component of movement trajectory, respectively; neither variable could explain ACC neural activity variance as well as sequence prevalence (Supplementary Fig. 9). Trial-by-trial values of expected reward, on the other hand, are challenging to estimate without explicit knowledge of the specific updating rule used by each animal. However, the robust performance of the prevalence model trained exclusively on ‘exploratory’ sequence instances (Supplementary Fig. 8a) argues against the possibility that changes in expected reward drive the observed activity modulation; in particular, our criterion for classifying a putative sequence instance as ‘exploratory’ inherently required that another behavioral sequence had locally become the animal’s dominant strategy, and only one sequence was ever eligible for reward in any task epoch. As such, most ‘exploratory’ sequence instances would not only have been unrewarded but would have likely been sampled by expert animals without any expectation of immediate reward collection. In further support of the conclusion that ACC neural dynamics reflect strategy prevalence rather than the associated reward expectation, we observed little activity modulation for sequence instances within ‘dominant’ tails – a period when sequence prevalence remains high, but reward expectation should rapidly diminish (Fig. 3f).

We also resolved whether the observed activity variation could simply reflect the specifics of choice and reward history that co-vary with prevalence arising from the pursuit of other strategies. To accomplish this, we evaluated whether the prevalence-based model trained only on a subset of sequence instances that shared immediate history still had significant explanatory power. Specifically, we capitalized on the previous finding that much of such direct history impact in the ACC is exerted within fewer than 5 trials (Bernacchia et al., 2011). Since the prevalence estimate is formed over a much longer window, we could sub-select those ‘LLR’ sequence instances that were matched in the immediate history (i.e. ‘LLR’ instances that immediately followed another ‘LLR’), but that otherwise still differed in the associated ‘LLR’ sequence prevalence. The prevalence-based model trained on this reduced subset of sequence instances matched for local history still explained a significant fraction of ACC activity variance (Supplementary Fig. 8b), arguing that the observed modulation could not have been solely history-based.

Overall, while our analyses do not rule out potential contributions of trial-by trial behavioral parameters to neural encoding in ACC, these parameters cannot account for the robustness of local strategy prevalence in explaining ACC neural activity variance. Combined, these data argue that strategy representation in ACC is continuously modulated by some summary statistic of that strategy in past choices, while large-scale ensemble reorganizations tag these representations with global contextual content.

### Strategy prevalence can be decoded from ACC ensemble trajectories during strategy execution

The unexpectedly strong modulation of ACC neural activity by sequence prevalence (Fig. 4) pointed to the possibility that information about how prevalent the currently sampled strategy has been in the recent past may be decodable from ACC ensemble activity by downstream circuits. Indeed, we found that linear population decoders trained independently in each analysis window achieved robust cross-validated performance. For each of the five analysis windows, we fit the relationship between the local sequence prevalence and the firing rates – in that window – of all simultaneously recorded ACC neurons. Cross-validated performance of such individually tuned linear decoders scaled steeply with the number of neurons (Supplementary Fig. 10). Indeed, ensembles of ACC neurons containing as few as 10 prevalence-modulated units could explain as much as 60% of sequence prevalence variance. Consistent with this finding, visualizing ensemble activity in a reduced dimensional state space defined from regression against prevalence (see Methods) showed clear separation of trajectories associated with different instances of sequence execution by the local strategy prevalence (Fig. 5a).

**Fig. 5.**
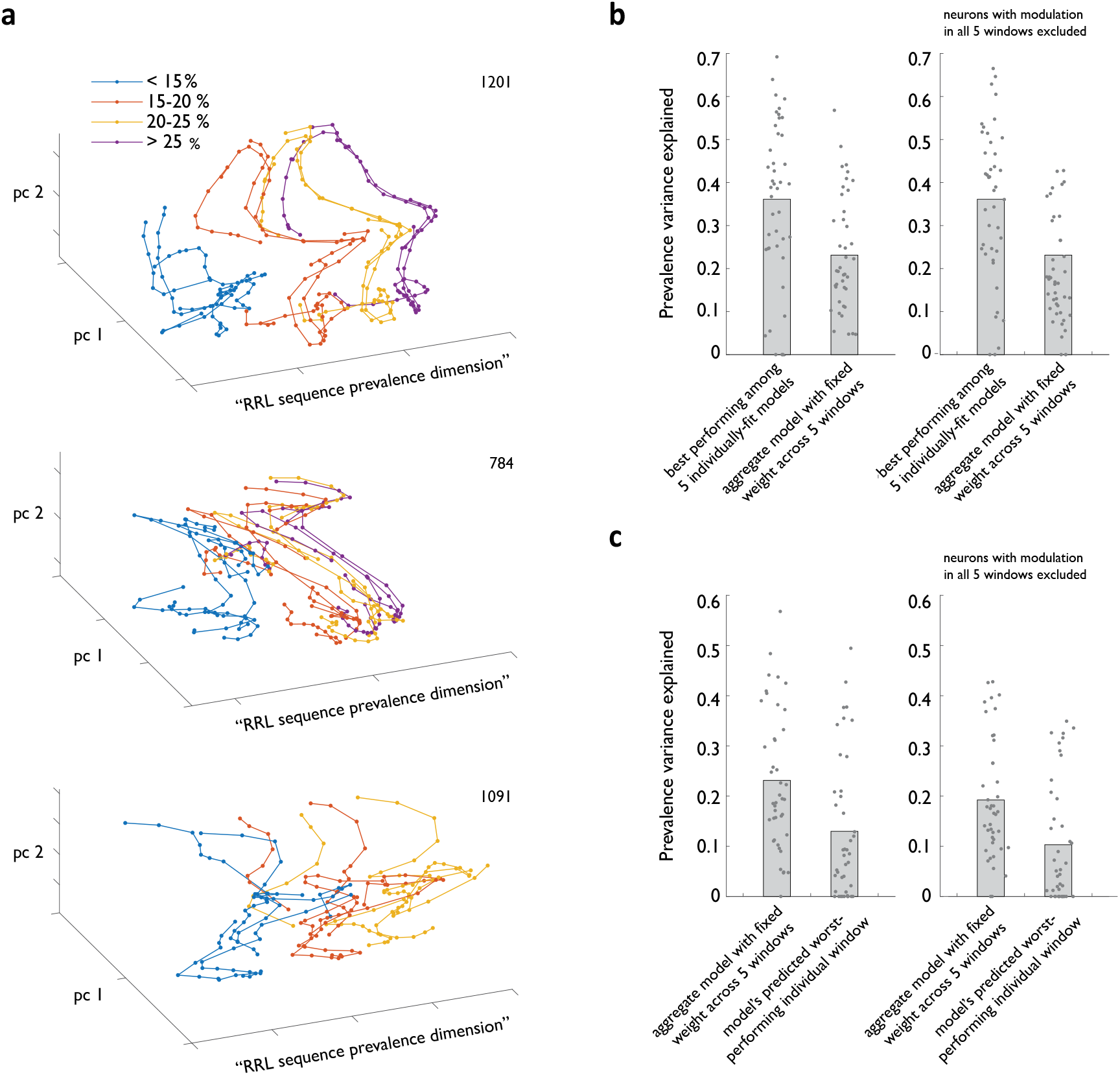
Strategy prevalence can be decoded from ACC ensemble activity throughout sequence execution. **a**. Examples from 3 different animals of ACC ensemble trajectories associated with different instances of RRL sequence execution visualized in ensemble state subspace chosen to maximize separation by local sequence prevalence. **b**. Cross-validated performance of linear models relating ACC ensemble activity during RRL execution to local RRL prevalence. Shown are the performance of best of 5 models fit in individual analysis windows **(b)**, the average performance of model that was constrained to always use the same weight for any given neuron in all five windows **(b, c)**, and that model’s worst predicted performance in an individual analysis window **(c)**. Right panels exclude from all models ACC neurons that show significant modulation by strategy prevalence in all 5 analysis windows.

The decoding strategy above implicitly assumes an ability to use a temporally varying readout, which may be challenging to implement in circuit dynamics. We therefore next examined the ability of temporally fixed readouts to decode prevalence. Focusing on sessions with at least 5 simultaneously recorded neurons that displayed modulation by prevalence in at least one analysis window, we next fit a new linear model that was constrained to always use the same readout (i.e. same weight for any given neuron in all five windows). The constrained model retained a large fraction of the overall explanatory power compared to that of the best of five individually tuned decoders (Fig. 5b, left panel, variance explained of 0.36 +/-0.03 for best of individually tuned models vs 0.23 +/-0.02 for the model with weights fixed across the 5 analysis windows). Moreover, when we evaluated the prediction of this aggregate constrained model in each of the five analysis windows, we observed substantial explanatory power even for the worst performing window (Fig. 5c, left panel), suggesting that the fixed decoder would never drop the prevalence signal as the animal executed the sub-steps of a multi-step strategy. The ability of a fixed linear sum of activity across the recorded population to retain a substantial fraction of explanatory power throughout the temporal extent of sequence execution was not simply due to the presence of a small number of neurons with prevalence-related modulation in all five analysis windows. Indeed, the constrained model’s performance dropped only moderately when such neurons were left out of the dataset (Fig. 5b-c, right panels). Combined, these observations suggest that despite the transient nature of activity modulation by sequence prevalence at the level of individual neurons (Supplementary Fig. 11), the ACC neural ensemble dynamics is structured in such a way as to permit this, or a related, summary statistic of the animal’s recent sequence choices to be stably decoded by downstream circuits throughout sequence execution.

## Discussion

The Anterior Cingulate Cortex is thought to play a central role in dynamic, context-specific strategy arbitration in complex non-stationary environments, yet the organizing principles of the cingulate’s neural activity that underpin contextually appropriate strategy selection and point to specific computations that take place in the ACC remain unresolved. We report here that when animals search a structured task space through self-guided exploration, a substantial fraction of ACC neural activity variance can be explained by the prevalence of individual behavioral strategies. This prevalence encoding – particularly enriched in the most rostral portion of the ACC homologous to the primate area 32D – is preserved through large-scale functional rearrangements of the ACC network between distinct global behavioral contexts and is evident even in the absence of any pairing between strategy execution and reward delivery. Our findings raise the possibility that the ‘attention to *self*-action’ (Blakemore et al., 2000) may be at the core of rostral cingulate functionality, in essence uniting the long-standing ‘attention to action’ account (Norman and Shallice, 1986; Passingham, 1996) with the proposed role of the cingulate in processing information relating to self (Blakemore et al., 2000; Euston et al., 2012; Passingham et al., 2010; Seamans and Floresco, 2021) and the production of self-generated actions.

Studies to define the precise role ACC plays in value-based decision-making have emphasized its role in tracking task-relevant information to guide appropriate action, but have otherwise often cast the question in terms of identifying the specific step to which it contributes in the pre-decision computation and comparison of action values (Boorman et al., 2009; Kennerley et al., 2009; Kolling et al., 2012; Luk and Wallis, 2013). The resulting lack of a unifying account has prompted a recent suggestion that rather than contributing to pre-decision valuation, ACC encodes post-decision variables related to the subjective value that can be inferred empirically from the animal’s choices (Blanchard and Hayden, 2014; Cai and Padoa-Schioppa, 2012). The notion of a subjective approach by each animal to the encountered task is similarly inherent in our parsing of the behavioral session using the animal’s selected self-guided strategy (rather than the imposed rule) in examining the encoding of behaviorally relevant information. Our observation that strategy prevalence explains a substantial fraction of variance in the activity of ACC neurons supports the interpretation that ACC computes an inherently subjective post-decision variable, and indeed, subjective value and strategy prevalence are closely related in well-trained animals in typical value-based tasks. However, our finding that prevalence contributes strongly to activity modulation even in the absence of reward delivery suggests that prevalence encoding might be a more parsimonious account that accommodates both past observations and our findings. Examining how adding such a summary statistic of the agent’s recent behavioral choices to models of decision-making changes their explanatory power in various experimental settings will be one interesting direction for future study.

The encoding of strategy prevalence appears to be independent of, and be preserved through, large-scale functional rearrangement of ACC networks. These functional rearrangements result in representational remapping, whereby ACC ensemble activity associated with a specific strategy changes markedly between behavioral contexts. This observation suggests that ‘network resets’ previously observed to accompany transitions to exploration (Durstewitz et al., 2010; Emberly and Jeremy, 2019; Karlsson et al., 2012) are not mere bookmarks of a behavioral state change, but rather reflect reorganizations of the ACC network designed to tag action plans with a global context-specific representation. Although ‘dominant’/ ‘exploratory’ distinction maps well onto the ‘default’/’alternative’ dichotomy that has enjoyed prominence in the ACC field due to its computational appeal and explanatory power in many settings (Blanchard and Hayden, 2014; Boorman et al., 2013; Kolling et al., 2012; Procyk et al., 2000), the representational transitions in ACC likely reflect broader contextual tagging of action plans, as suggested by the observed difference in the representations associated with a specific exploratory sequence depending on the nature of the default strategy. It is even possible that the contextual content of the neural representations in ACC may not be limited to the statistics of actions and outcomes. Indeed, given the ACC’s topological centrality within the frontal cortical network and domain-general role in cognition, the content of its neural representations set-up by large scale re-arrangements of functional networks might come to reflect, when appropriate, distinct cognitive loads (Shenhav et al., 2016), social settings (Tomlin et al., 2006) or somato-visceral states (Caracheo et al., 2018; Seamans and Floresco, 2021). As such, ACC would set up – and possibly initially infer — representations of distinct task-relevant contextual information in such a way that tracking of individual strategy prevalence may then take place.

How is the circuit implementation of the abrupt representational re-organizations that define individual behavioral contexts related to prevalence encoding? One possibility is that these representational re-organizations are simply an emergent property produced by the dynamic interactions among the elements of the network that tracks prevalence. Specifically, a gradual, prevalence-tracking change in the activity of individual neurons could eventually reach a threshold that instantiates a sudden phase transition reorganizing ACC into a new functional network. Against this idea, prevalence often changes abruptly rather than slowly at the end of a behavioral block, and furthermore, changes in prevalence inside ‘dominant’ blocks often exceed those at the end of a behavioral block but do not lead to abrupt reorganizations of the network. Alternatively, the ACC network may be built to allow a continuous representation of prevalence through a substantial degree of reorganization and a different computation triggers the abrupt transitions. What constraints such an encoding scheme places on the ACC network architecture, where the transition-triggering computation is performed, and what factors into that computation remain open questions. Furthermore, investigating what constraints an encoding scheme that preserves prevalence computation through large-scale functional rearrangements places on network architecture and biophysics may likewise be an intriguing area of future study.

What might be the computational advantage of encoding prevalence, and how can the encoding of this post-decision variable be reconciled with the recent evidence causally linking ACC to the current decision (Tervo et al., 2021)? Given ACC’s well-documented role in exploration (Blanchard and Hayden, 2014; Fouragnan et al., 2019; Hayden et al., 2011; Kolling et al., 2012; Quilodran et al., 2008; Tervo et al., 2021), one possibility is that keeping track of strategy prevalence may be used to prioritize exploring strategies evaluated less frequently in the recent past (see (Wiering and urgen Schmidhuber) for one implementation of such frequency-based exploration). An alternative view is informed by old accounts of the ACC’s role in recognizing the ‘self’ actions (Blakemore et al., 1998; Espinosa et al., 2006). Indeed, an intriguing possibility is that in complex settings, keeping track of different actions taken and the frequency with which one has performed them might be an effective way of estimating agency – an understanding of which actions bring about the ability to promote or prevent the occurrence of events in the environment – through statistical learning. As such, agents can determine the degree to which their actions exert control over world events without reinforcement or an explicit representation of the temporal relationships between their actions and events. Statistical learning is thought to be central to many aspects of perceptual cognition and motor control; in the causal domain, it provides a complement to associative learning in permitting the agent to make inferences about the world without being enslaved to temporal contiguity or specific action-outcome contingencies. In principle, learning of statistical regularities can happen implicitly – without explicit awareness or hypothesis-testing. However, one influential view posits that behind the efficiency, with which animals pick up on statistical regularities from sparse data in complex, often only partially observable environments, is the process of probabilistic inference that entertains multiple candidate models of the underlying statistical regularities coupled with explicit hypothesis testing designed to evaluate those models (Tenenbaum et al., 2011). And indeed, establishing agency would critically depend on active evaluation of an inferred ability to influence events in the world. We posit that ACC’s central role in establishing one’s agency is what reconciles our finding of its robust prevalence encoding and the causal evidence (Tervo et al., 2020) linking it to the ongoing decision of whether or not to evaluate alternative action plans (and more generally to curiosity (Wang and Hayden, 2020), information seeking (White et al., 2019) and hypothesis testing (Elliott and Dolan, 1998)).

We note that the interpretation that ACC – and likely the medial frontal network into which it is embedded – plays a key role in evaluating the extent to which one has agency in the environment is a refinement of the broader hypothesis that it processes information related to self and implements self-generated actions that not only accommodates the existing experimental observations but is also more falsifiable. Indeed, a major critique of the ‘self-generated actions’ view of the medial frontal lobe is that the distinction between self-generated and externally-triggered actions is empirically intractable, thus making this view of the medial frontal lobe unfalsifiable (Schüür and Haggard, 2011). In contrast, it is conceivable to develop experimental designs that manipulate the reward statistics to induce miscalculations of agency: superstitions – a perception of control over the environment in the absence of a causal link between the agents’ actions and the outcome and learned helplessness – an incorrect perception of a lack of control and a cessation of action.

## Acknowledgements

We thank Elena Kuleshova for help with surgeries and histology. We thank Shaul Druckmann, Gowan Tervo, Michael Brainard and Vivek Jayaraman for advice and comments on the manuscript.

## Author Contributions

M.P., M.M. and A.Y.K. designed the behavioral training protocol and trained animals to perform the task. M.P and M.M. performed the electrophysiology experiments and analyzed the data. M.P., M.M. and A.Y.K. wrote the manuscript.

## Data and materials availability

Requests for data, code and materials should be addressed to A.Y.K.

## Supplementary Figure Legends

**Supplementary Fig. 1.**
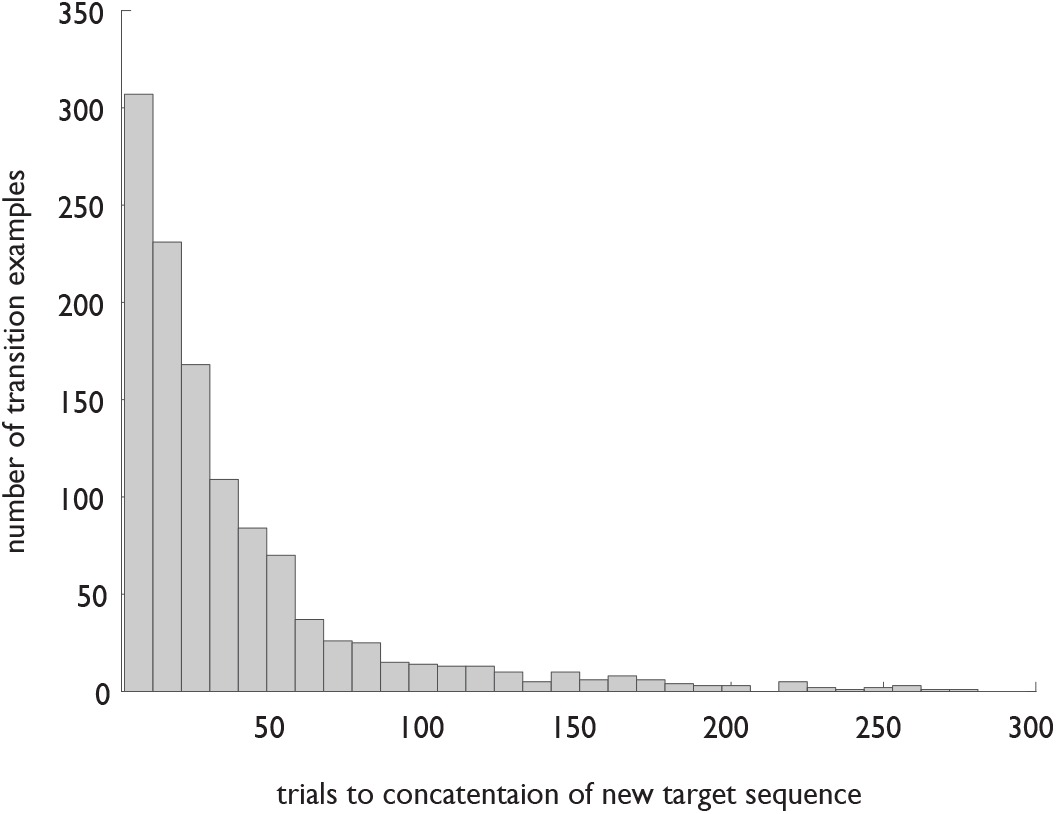
Rats efficiently adjust to unsignalled changes in the identity of the latent target sequence. Distribution of trials to concatenation of the new target sequence for all transitions in the dataset from the implanted animals.

**Supplementary Fig. 2.**
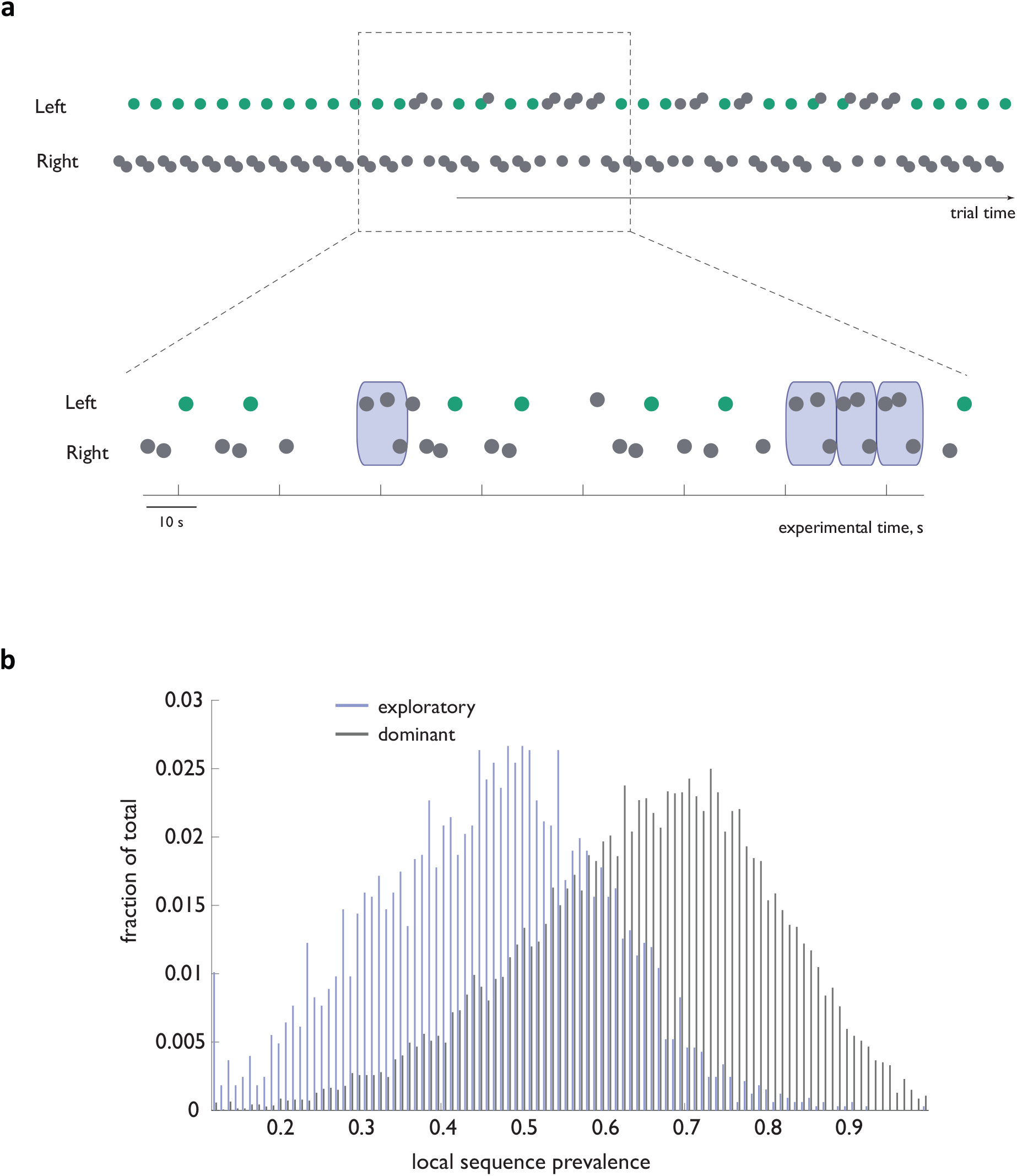
Exploratory deviations from the dominant sequence occur even in the absence of reward omission. **a**. Example behavioral trace from an animal that never experienced reward omission (ie the dominant sequence was always rewarded with 100% reliability) highlighting a mid-block transient exploratory bout. **b**. Distribution of local sequence prevalence values for the ‘Left-Left-Right’ and ‘Right-Right-Left’ sequences across all dominant and exploratory instances in the dataset from the subset of the implanted animals that never experienced reward omission.

**Supplementary Fig. 3.**
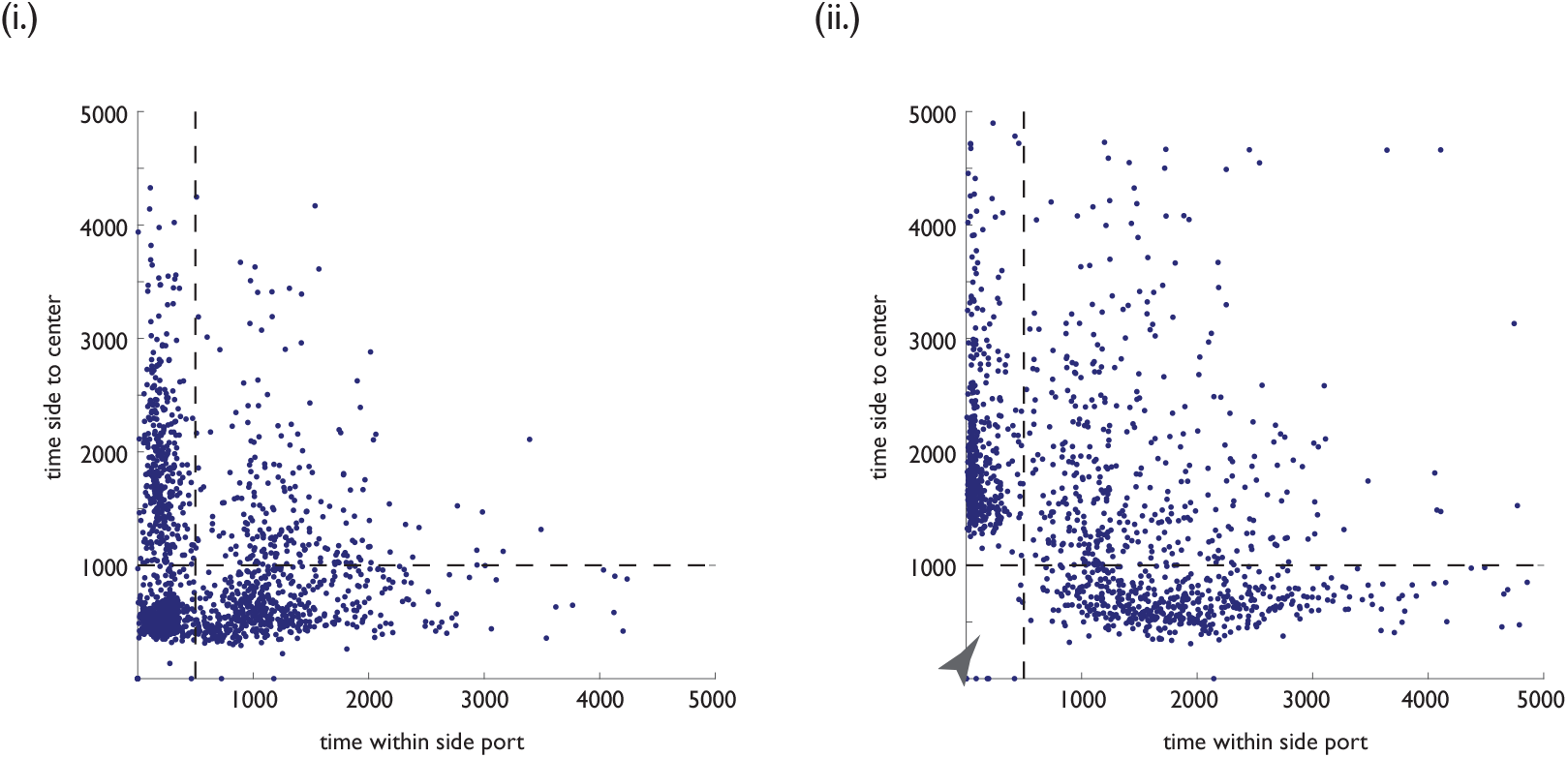
Delayed exit from the side port on the last step of unrewarded legitimate sequence instances aids in exploratory sequence classification. Side-to-center time vs within-side-port time distribution for all dominant sequence instances, for which reward was omitted. **(i):** steps one in the sequence. **(ii):** step three in the sequence. Note the conspicuous absence of short durations for the third sequence step (arrowhead).

**Supplementary Fig. 4.**
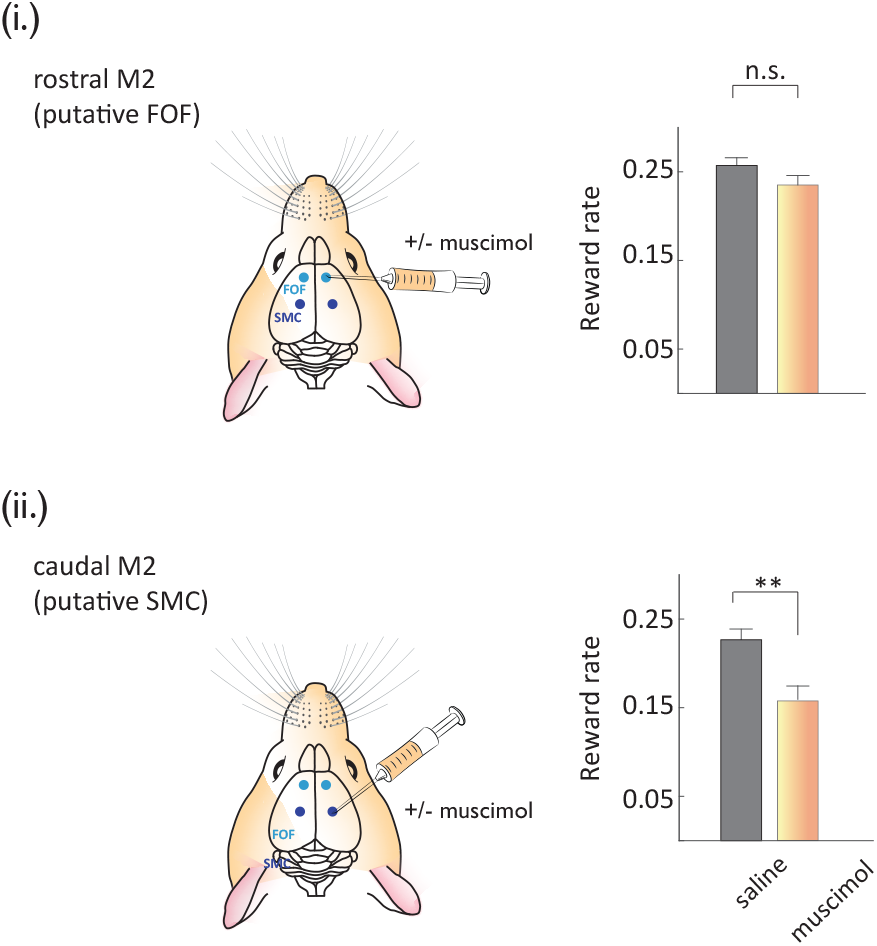
Pharmacological inactivation of a putative rat homologue of the Supplementary Motor Cortex (SMC) impairs self-guided higher order action sequencing. **Left panels:** schematic of muscimol delivery to the rostral part of agranular secondary motor cortex M2, putatively FOF **(i)**, or posterior part of M2, putatively SMC **(ii). Right panels:** Performance on the self-guided sequence task (as average reward rate) during saline and muscimol injections in the target region. Animals sample the ‘non-preferred’ option irrespective of the relative reward rate. Note that the basic sequencing of actions pairing initiation port and side port entries to complete a trial is not impaired when the putative rat SMC is inactivated.

**Supplementary Figure 5.**
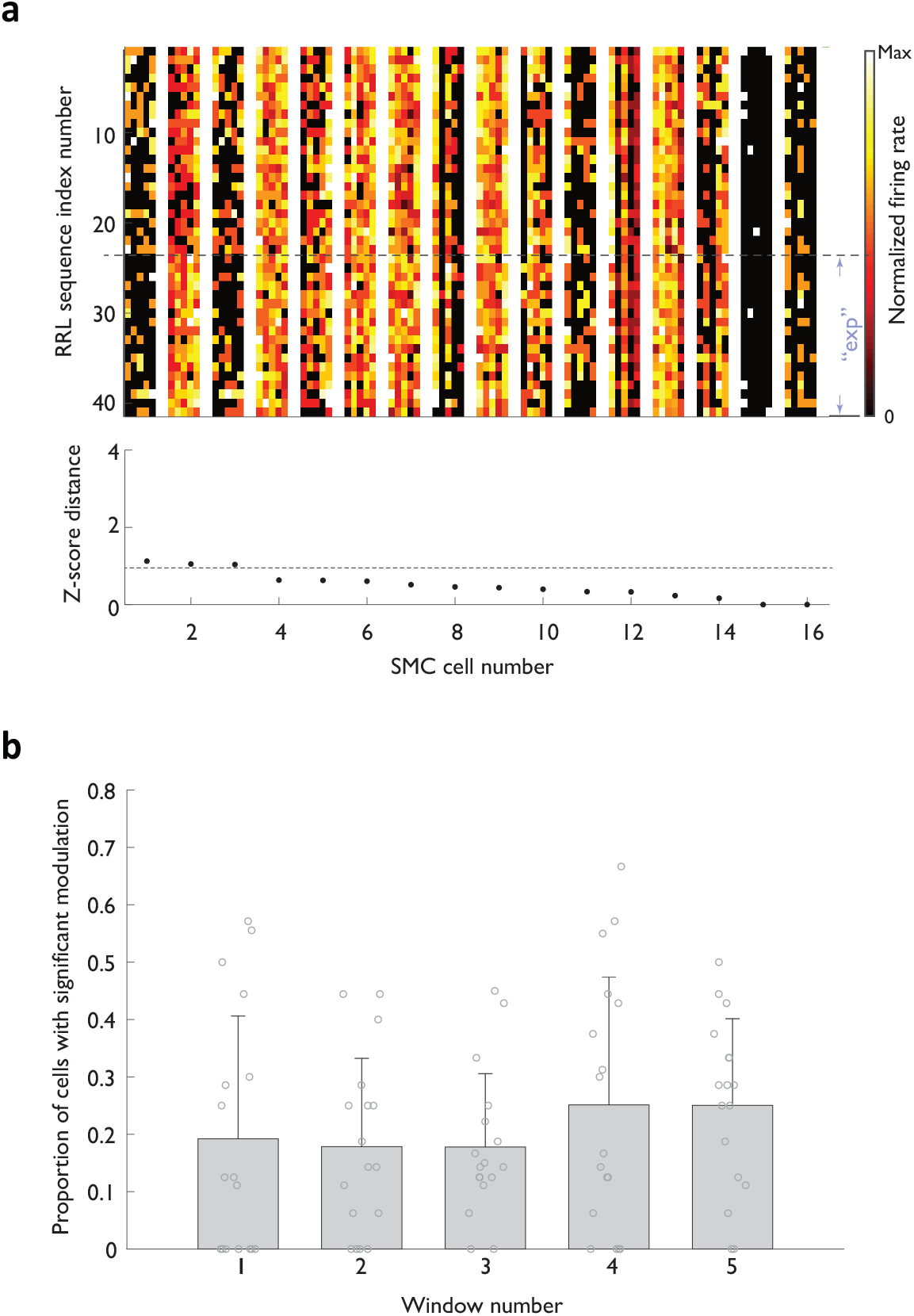
Representational transitions are less prominent in the caudal portion of the medial frontal lobe. **a**. Heat map representations of normalized activity associated with ‘Right-Right-Left’ sequence execution for 16 simultaneously recorded SMC neurons. Activity profiles across different sequence instances are stacked vertically. ‘exp’-‘exploratory’ instances. Neurons are arranged according to a ‘transition score’ defined as the distance between the two cloud centroids normalized by root mean of variance within each cloud. Transition score range reflected on the plot was chosen to match that in Figure 3b. **b**. Fraction of SMC neurons displaying significant context-related transition in each of the five analysis windows.

**Supplementary Figure 6.**
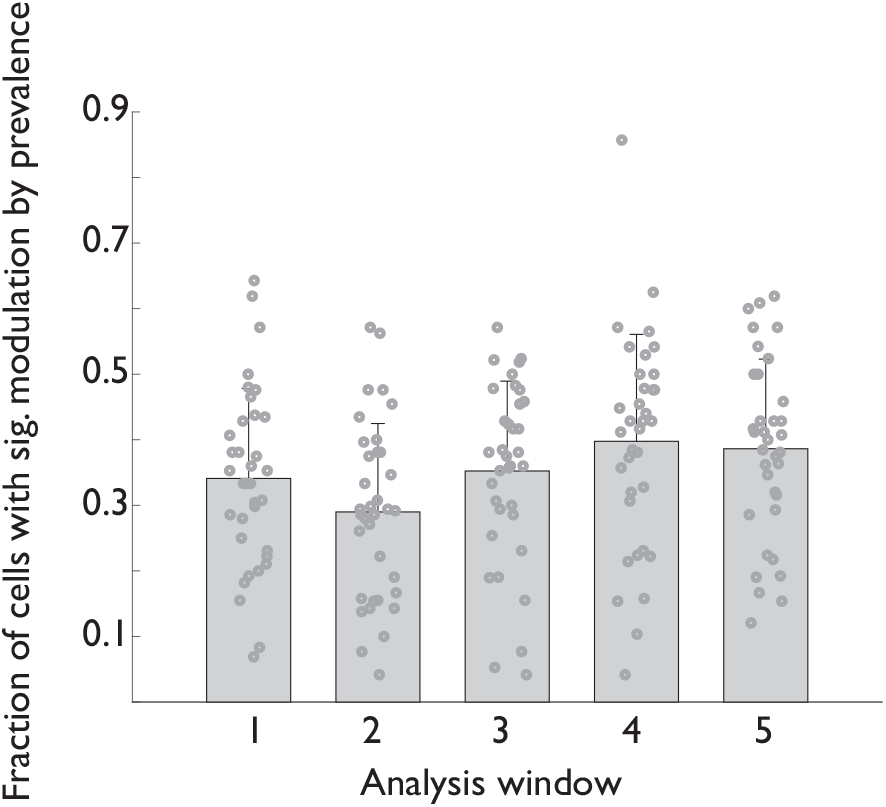
Modulation of ACC activity by strategy prevalence is evident in all 5 analysis windows. Fraction of all recorded ACC units displaying a significant modulation by sequence prevalence for each of the five analysis windows.

**Supplementary Figure 7.**
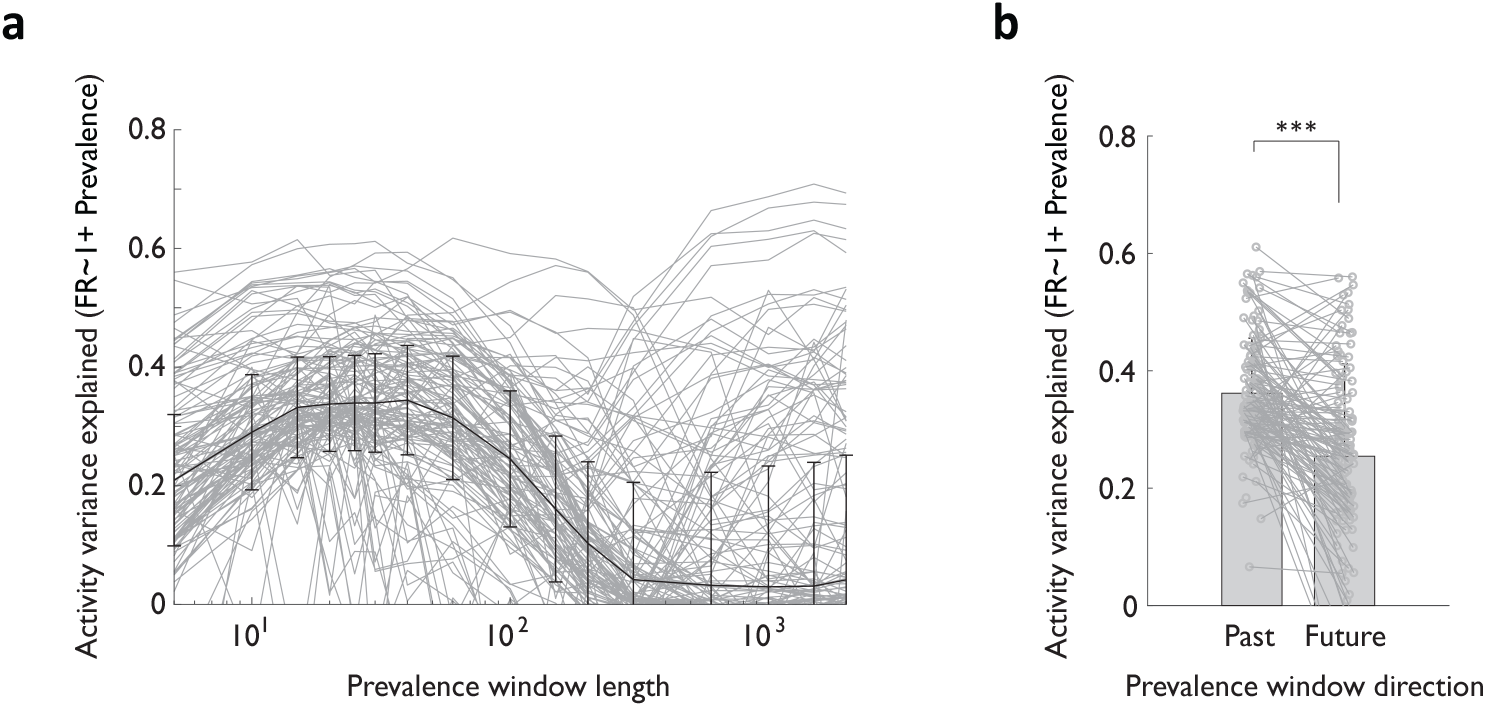
Models relating ACC neural activity to strategy prevalence in recent (within tens of trials) past display the strongest explanatory power. **a**. Explained variance in neural activity associated with RRL sequence execution – for models without context as a parameter – as a function of the number of trials in recent past used to estimate local RRL sequence prevalence. **b**. Explained variance in neural activity associated with RRL sequence execution for models that use local sequence prevalence in recent past (Past) or in the equivalent upcoming period (Future).

**Supplementary Figure 8.**
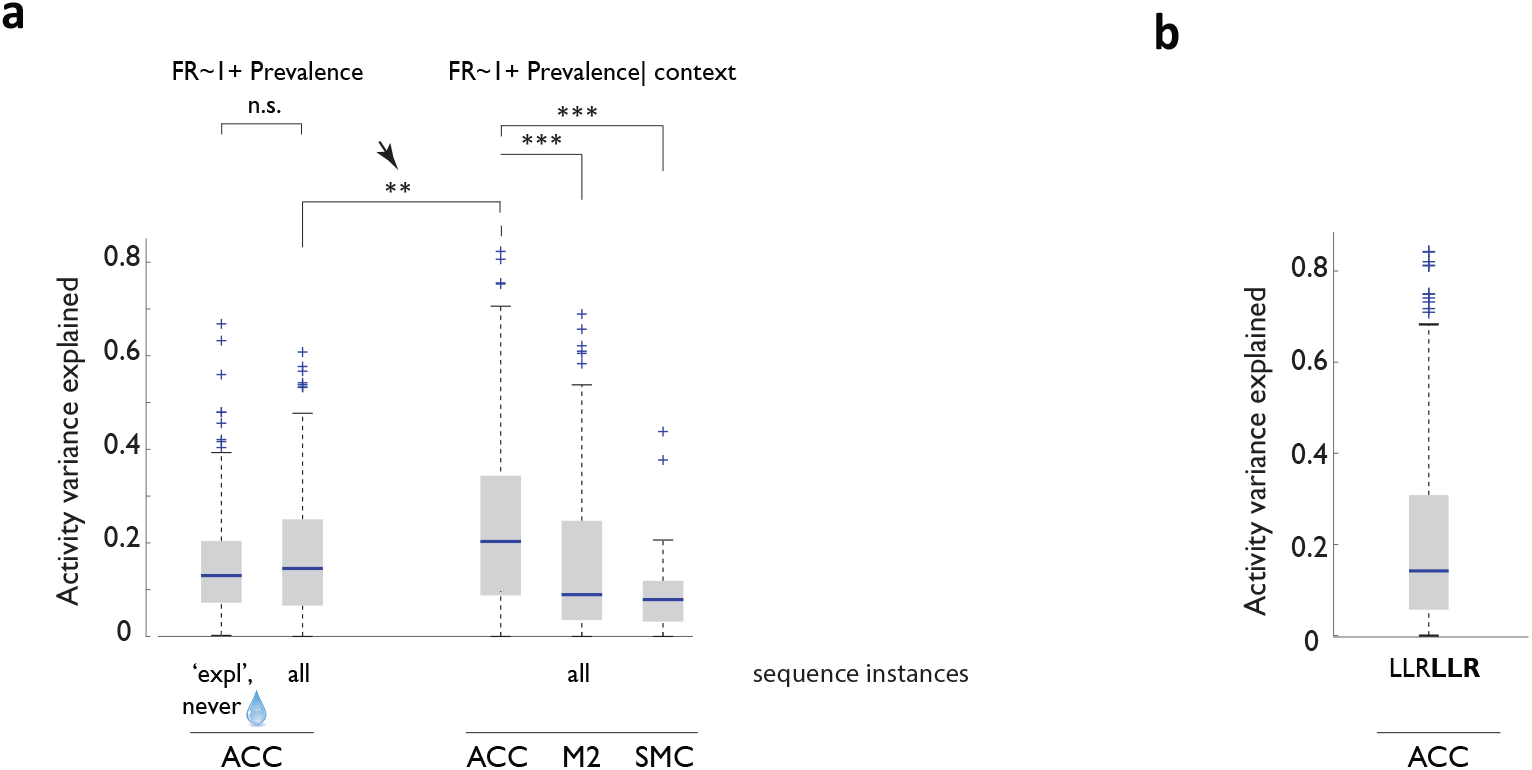
Models relating neural activity to strategy prevalence retain robustness when trained on subsampled data that controls for history of choices and rewards and display more robust performance in ACC than in other areas of the medial frontal lobe. **a**. Cross-validated performance of linear models relating local sequence prevalence (with and without global context – ‘exploratory’ or ‘dominant’ as a fixed parameter) to ACC, M2 and SMC neural activity. *exp*, fit done for exploratory sequence instances only; *all*, fits done for all sequence instances. Arrow points to improved performance when global context is included as a parameter. **b**. Cross-validated performance of the prevalence model trained on a subsample of the dataset matched in recent history.

**Supplementary Figure 9.**
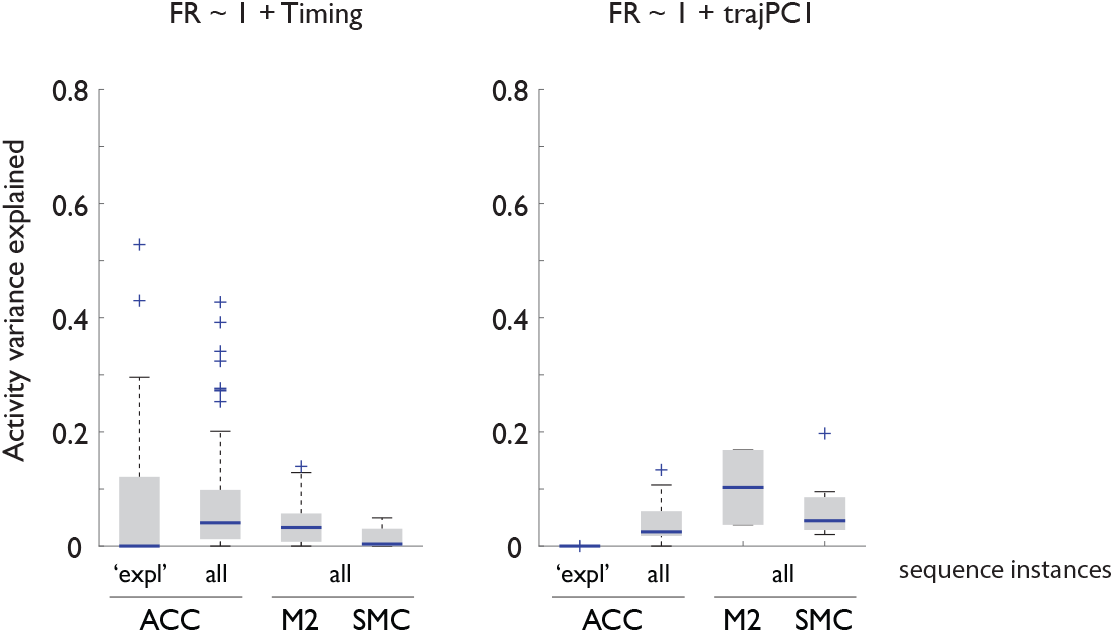
>Variance of movement vigor and trajectory cannot account for the robustness of sequence prevalence in explaining ACC neural activity variance. Cross-validated performance of linear models relating sequence time (left) or the first principal component of the associated trajectory (right) to ACC, M2 and SMC neural activity.

**Supplementary Figure 10.**
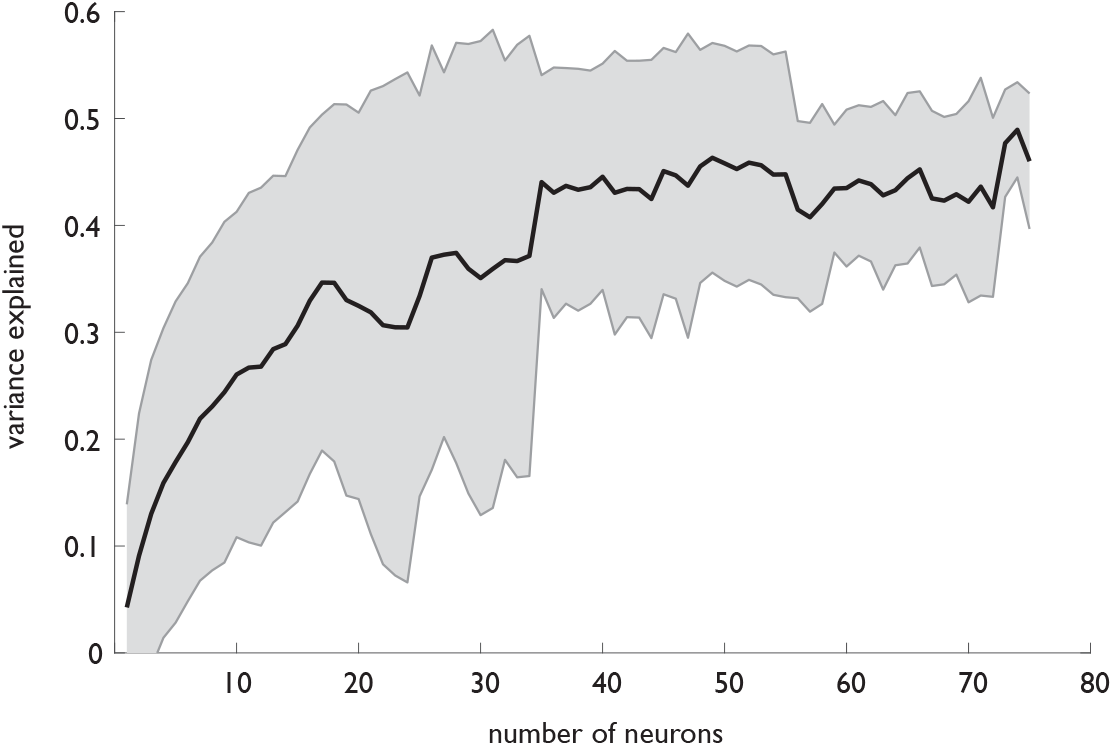
Prevalence decodability scales with the ACC ensemble size. Explained variance of prevalence (estimated using the 20-trial window of past choices) as a function of the size of the ACC ensemble.

**Supplementary Figure 11.**
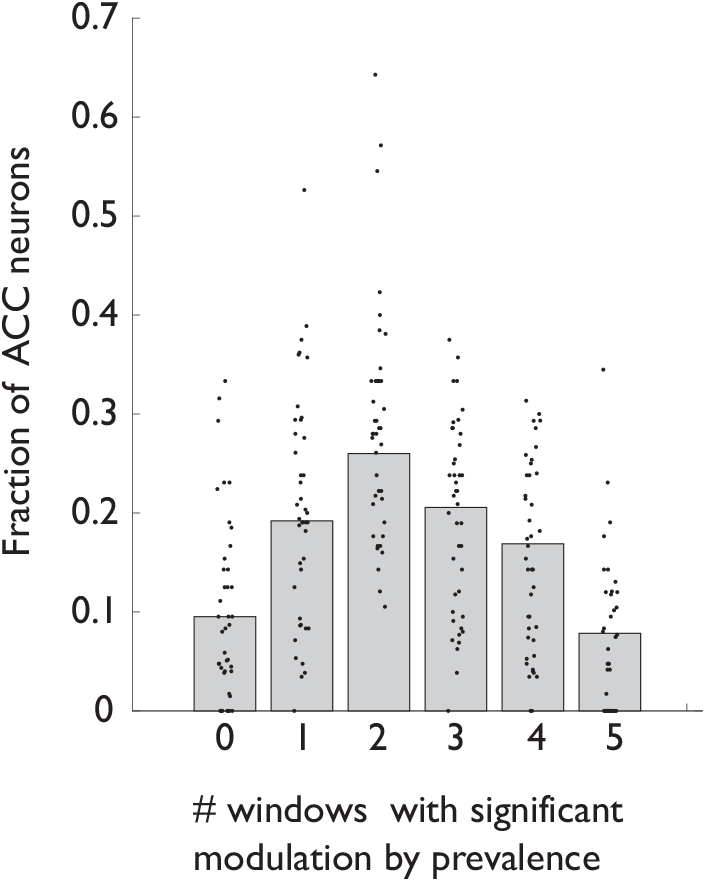
At the single neuron level, modulation by strategy prevalence rarely persists throughout the temporal extent of strategy execution. Fraction of ACC neurons displaying modulation in a specific subset of the five analysis windows.

## Methods

### Subjects

All experiments were done in male Long Evans rats 6-12 months of age (with weight kept between 400 and 500g). Animals were kept at 85% of their initial body weight before food restriction and maintained on a 12hr light/12hr dark schedule. Experiments were conducted according to National Institutes of Health guidelines for animal research and were approved by the Institutional Animal Care and Use Committee at HHMI’s Janelia Farm Research Campus.

A total of 7 animals were implanted with tetrode drives for collection of neural activity during the task (4 animals targeting the ACC and 3 animals targeting the SMC).

### Behavioral apparatus and task

All behavior was confined to a box with 23 cm high plastic walls and stainless-steel floors (Island Motion Corp). The floor of the box was 25 cm by 34 cm, and the nose ports were all arranged on one of the 25 cm walls. All lights, nose ports, and reward deliveries were controlled and monitored with a custom-programmed microcontroller, which in turn communicated via USB to a PC running a Matlab-based control program. Nose port entries were detected with an infrared beam-break detector (IR LED and photodiode pair). The central initiation port contained one white LED that indicated the option to initiate a new trial. The side choice nose ports also each contained an LED that indicated that the initiation port had been successfully triggered (at which point the LED in the center port was extinguished) and side ports were available for selection. Note that in some sessions, only center LED was changing states, and in some no lights were used with little impact on behavioral performance. The side ports also delivered liquid rewards (0.1 ml drops of 10% sucrose mixed with black cherry Kool-Aid) with the help of a motorized syringe pump (Harvard Apparatus PHD 2000).

The behavioral task is an elaboration of the basic design reported as a ‘covert pattern’ task in (Tervo et al., 2014) and consists of a series of self-initiated trials (several hundred per session), each involving a choice between two options – the left and the right choice port. Reward is delivered on the last step of a target sequence that is not otherwise indicated to an animal and thus has to be discovered through self-guided exploration.

Each session in the dataset described in this manuscript contained one to several unsignalled transitions in the identity of the target sequence. The set of possible sequences typically included some or all of the four non-trivial three step sequences (‘Left-Left-Right’, ‘Right-Right-Left’, ‘Left-Right-Right’, ‘Right-Left-Left’), but occasionally was expanded to include longer sequences (mainly ‘Left-Left-Left-Right’). Only one sequence was rewarded in any particular block of trials. A subset of sessions incorporated reward omission for 10-30% of correctly executed sequences.

### Behavioral training

Food-restricted animals were trained to perform the task with no explicit instruction. Early in training, exploration was encouraged by rewarding a small fraction (at most 10%) of novel patterns – specifically, those that escaped prediction by Competitor 2 (Tervo et al., 2014) that used the history of the animal’s choices and outcomes to predict his next choice. Note that once the animal discovers and concatenates the target sequence at a high rate, little to no background reward is delivered since the behavior becomes fully predictable. Eventually, background reward was fully eliminated.

Animals were considered proficient on the task when they consistently discovered and concatenated more than one sequence in a session, and when the average across-session reward rate was consistently in the 17-22 % range % (with 33% being the maximal theoretically possible). Over 90% of animals became proficient within one month of training.

### Electrophysiological recordings

A microdrive array containing 16 independently movable tetrodes was chronically implanted on the head of the animal. Each tetrode was constructed by twisting and fusing together four insulated 13 μm wires (stablohm 800A, California Fine Wire). Each tetrode tip was gold-plated to reduce impedance to 200-300 kΩ at 1 kHz. Within the implant, the tetrodes converged to an oval bundle (1 mm x 2 mmd), angled at 0° with respect to vertical (pointing towards midline after implantation).

For the drive implantation surgery, trained animals were initially anaesthetized with 5% isoflurane gas (1.0 L/min). After 10-15 minutes, isoflurane was reduced to 1.5-2.0% and the flow rate to 0.7 L/min. A local anesthetic (Bupivacaine) was injected under the skin 10 minutes before making an incision. A unilateral craniotomy (1.0 by 2.0 mm) was drilled in the skull above the site of recording. The microdrive array was implanted such that the tetrode bundle was centered 3.0 mm anterior and 0.8 mm lateral to Bregma (right or left hemisphere) when recordings were targeted to the ACC, and 2.0 mm posterior and 1.2 mm lateral to Bregma when recordings were targeted to the putative SMC. Small stainless steel bone screws and dental cement were used to secure the implant to the skull. One of the screws was connected to a wire leading to the system ground. Before the animal woke up, all tetrodes were advanced into the brain ∼1.20 mm deep from the brain surface.

Over the two weeks following surgery, the tetrodes were slowly lowered, moving approximately 40 μm/day on average. During this time, animals were re-acclimated to performing the task with the drive. When performance on the task was regained to pre-surgery levels (in terms of motivation and dynamic strategy arbitration behavior), recording sessions began. After each recording session, any tetrodes that did not appear to have any isolatable units were moved down 25 μm. Once a tetrode had been moved a total of 2.5 mm from the surface, which is the approximate border between anterior cingulate and prelimbic cortices, it was no longer advanced.

Each recording session spanned 3 to 4 hours. Animals were not forced to perform the task and sometimes took breaks (generally around 10 minutes, but sometimes up to 30 minutes). Data from all the animals were collected using the wireless headstage and datalogger (Horizontal Headstage 128ch with Datalogger, SpikeGadgets, spikegadgets.com/hardware/hh128.html). An array of LEDs of different colors was attached on top of the animal’s implant and the animal’s position in the environment was recorded with video camera at 60 frames per second. The animal’s position was reconstructed offline using a semi-automated analysis of digital video of the experiment with custom-written software.

Raw electrophysiology data were sampled at 30 kHz, digitally filtered between 600 Hz and 6 KHz (2 pole Bessel for high and low pass) and threshold crossing events were selected for further analysis. Individual units on each tetrode were identified by manually classifying spikes using polygons in two-dimensional views of waveform parameters using custom made Matlab scripts (Karlsson et al. 2012, MatClust, https://www.mathworks.com/matlabcentral/fileexchange/39663-matclust). For each channel of a tetrode, peak waveform amplitude and the waveform’s projection onto the first two principal components were used for clustering. Autocorrelation analyses were done to exclude units with non-physiological single-unit spike trains. Only units where the entire cluster was visible throughout the recording session were included. Thus, a unit was not isolated for further analysis if any part of the cluster vanished into the noise or was cut off by the recording threshold.

The total contribution from each animal was as follows:

ACC animals: 4 sessions, 81 neurons for animal 1, 15 sessions, 163 neurons for animal 2, 13 sessions, 324 neurons for animal 3, 5 sessions, 286 neurons for animal 4

SMA animals: 11 sessions, 176 neurons for animal 1, 12 sessions, 101 neurons for animal 2, 8 sessions, 75 neurons for animal 3

### Mapping of the putative SMC homologue

The Supplementary Motor Cortex had not been characterized in the rat, so in a separate set of experiments, we sought to identify a medial premotor region involved in temporally sequencing self-initiated actions in the rat. Specifically, we performed a set of pharmacological inactivation experiments along the rostro-caudal axis of the agranular premotor region M2, evaluating the effect of such a transient inactivation on the animals’ performance during the three-step version of the sequence exploration task. We observed a robust behavioral effect following a bilateral injection of muscimol, but only when muscimol was delivered to the region immediately caudal to mid-cingulate cortex (a location similar to that of primate SMC, Supplementary Fig. 4). Following local muscimol delivery, animals continued to complete trials, but no longer performed the target sequence significantly above the value expected for a biased coin. SMC inactivation appeared to specifically affect complex sequencing rather than chaining of actions in general, because the animals still performed the sequential entries into the initiation and reward ports correctly. The technical details associated with this set of experiments are provided below.

### Surgery for Cannula Implantation

Location of bilateral craniotomies and cannula implantation were +2.0 mm AP and ± 1.2 mm ML with respect to Bregma for ACC, -2 mm AP and ± 1.2 mm ML for putative SMC, +1mm AP and ± 1.2 mm ML for FOF (Erlich et al., 2011), -0.5 mm AP and ± 1.2 mm ML for another candidate SMC region that didn’t show muscimol effect. ACC canula implantation was deeper (2.0-3.)0 mm to account for curvature, whereas candidate SMC regions were 1 mm deep. Injection guide cannulas (Eicom CXG (T) 2 Diameter OD/ID 0.3/0.2 mm) were inserted through a 0.5mm craniotomy. As a protection measure for the animal in the home cage, an opaque cone was placed around the implant. Cannulas were bonded to the skull with dental cement (C&B Metabond -Parkell). Dummy cannulas (Eicom CXD (T) 2 Diameter 0.15/0.06 mm) were used in between sessions to protect the cannula from debris. Food deprivation and further training was resumed after recovery period.

### Muscimol Inactivation

As a control, the beginning of each session remained unperturbed and animals were allowed to perform the task normally. After an animal performed 200 trials and reached reward rate above 0.22, the animal was placed back into its home cage for inactivation. Treats were provided to keep the animal still. Muscimol or saline was administered through the needle inserted into cannula. Muscimol (Tocris Bioscience) solution was prepared with sterile saline (Hospira) at a final concentration of 0.1 μg/ml. Bilateral infusion was made via a 3 mm-long 31-gauge injector connected to a 5 μl syringe (Hamilton) by a teflon tube (Eicom). Eicom micro syringe pump ESP-64 was programmed to deliver the solution at the rate of 0.25 μl/min for 2 min, for a total volume of 0.50 μl for each hemisphere. The animal was placed back into the operant chamber after the injection. Muscimol effect persisted for about two hours after the injection. Trials within one hour of the injection were used for the analysis.

### Data analysis

#### Session selection

Since animals frequently showed greater variability in behavioral performance following drive implantation, a minimal selection filter was applied to determine, which sessions to include into the analysis: the session had to contain at least 200 instances of either the ‘LLR’ or the ‘RRL’ sequence, and the across-session average sequence production rate had to exceed 0.15.

#### Selection of dominant and exploratory sequence instances in behavioral data

The nature of our behavioral framework required extra care when parsing the behavioral record to identify ‘legitimate’ sequence instances. Parsing a continuous stream of left and right choices to identify ‘legitimate’ sequences was easy for the ‘dominant’ condition, where a high prevalence of sequence concatenation, and a scarcity of choices that conform to other patterns, argue that almost every instance of a pattern conforming to the target sequence is likely to be one actually evaluated by the animal. Specifically, starting with all ‘LLR’ (‘RRL’) instances that were done during the epochs where ‘LLR’ (‘RRL’) was the latent target sequence, we removed all examples prior to when the dominance for the new sequence (which we define as the presence of a successful concatenation of at least 3 sequence instances) hadn’t yet been established, and appended all sequence instances from the period where dominance had persisted following a block change (i.e. until the previously concatenated sequence was interrupted by more than two trials incompatible with that sequence).

The ‘exploratory’ condition, required a closer examination outside of clear concatenations. For example, although some ‘RRL’ patterns within runs of ‘LLRRL’ in a ‘LLR’ block might reflect a true pairing of the locally dominant pursuit of ‘LLR’ with a quick exploratory evaluation of ‘RRL’ tagged on in a manner akin to strategy mixing (Donoso et al., 2014), most of such instances likely reflect a mere apposition of the ‘RRL’ and the ‘LR’ sequences. To select the likely true exploratory sequences conservatively, we therefore excluded all putative lone exploratory sequences that displayed an overlap (to the left of to the right) with the dominant sequence, *except* when the timing analysis (see below) provided independent evidence in favor of classifying this putative instance as a true exploratory sequence. Here, our approach was grounded in the expectation that animals would pause, if only briefly, at the side nose port on the last step of a true exploratory sequence instance to ascertain that the explored sequence is not rewarded. Under this assumption, we ‘rescued’ the putative exploratory sequence instances for which the duration of the third step exceeded the threshold chosen on the basis of an unbiased analysis of the distributions for the within-sideport and side-to-center durations across all ‘dominant’ sequence instances for which the otherwise scheduled reward was omitted permitted an unbiased selection of the specific threshold (Supplementary Fig. 3). Specifically, starting with all putative ‘LLR’ (‘RRL’) during periods when dominance had been established for another sequence (see above), we first removed all instances that displayed an overlap (to the left, or to the right) with the dominant latent target sequence. From the removed set, we then ‘rescued’ those instances that were either preceded by another ‘LLR’ (‘RRL’), or for which on step 3, the within-sideport duration exceeded 0.5 seconds, or side-to-center duration exceeded 1 second.

In addition, we noticed that at times, animals chose to concatenate ‘LLRR’, possibly as a clever way to catch every three step sequence that might serve as the latent target. Therefore, we removed from all datasets any ‘LLR’ (‘RRL’) instances that appeared within such concatenations. Finally, we also removed from all datasets sequence instances where the animal appeared to take long breaks. Specifically, any sequence instance that displayed a center-to-side time of over three seconds was removed from further analysis.

#### Analysis of sequence representation in frontal cortical activity

To capture the neural activity associated with a specific three-step sequence, we represented, for each neuron, its activity during a single instance of sequence execution as a vector of the square root of spike counts in five 500 millisecond windows centered on port entry events (center and side port entries for steps 2 and 3 in the sequence, and side entry for step 1; the window leading to center port entry in step 1 was excluded to avoid a strong contribution of behavioral choices that preceded sequence initiation). The choice of working with the square root of spike counts was guided by the desire to transform the data with Poisson distribution into an approximately Gaussian distribution with a unity variance (Anscombe, 1948). Such transformation brought the variance of the binned data for individual cells to the same level, such that the variability of fast spiking neurons did not conceal less active cells. The five chosen windows offered substantial coverage of the sequence with minimal temporal overlap, a high degree of stereotypy of spatial trajectories, and the exclusion of outcome-related activity.

For a single neuron, its activity associated with each instance of sequence execution was thus captured as a single point in the corresponding five-dimensional ‘state space’. A single-cell metric of the contextual representational change (‘transition score’, Figure 3) was calculated as the Euclidean distance in this single cell state space between the centroids of the ‘dominant’ and the ‘exploratory’ clusters.

#### Decoding analysis

For each session, we trained a linear classifier to predict, on the basis of the ensemble activity, whether a particular trial was a part of the ‘LLR’ or the ‘RRL’ sequence. For each trial, we used firing rates of simultaneously recorded ACC cells as predictors and sequence identity as a predicted variable. All cells with non-zero firing rate were included in the analysis; firing rate for each was calculated over the 500 msec decoding window centered around entry into the side port. When the entire 3-step sequence identity was being decoded, each cell contributed 3 windows corresponding to the 3 side entries. When the decoding was done on the context (‘LLR’ vs ‘RRL’) of a specific choice (e.g. ‘Right’), only one window was used. All linear discriminant analyses were performed with a 5-fold cross validation: 5 separate classifiers were trained on 80% of the data set, and the classification accuracy for each was estimated on the remaining 20% of the dataset, after ensuring that the sample size in each category was the same. The reported values represent the averaged the error rates across the 5 classification runs.

#### Characterization of representational transitions

To capture the behavior of the entire neuronal ensemble, the dimensionality of the state space was expended to 5*N, where N was the number of simultaneously recorded cells. We then calculated pairwise Euclidian distance between the points in the state space to evaluate how the sequence representation evolved over the course of the session. To demonstrate clustering of points according to the behavioral context, we calculated the distance between centroids of the clusters (median values along each dimension) corresponding to the ‘dominant’ and ‘exploratory’ groupings, and compared it with the corresponding distance calculated after context identity was randomly shuffled across the differences sequence instances.

#### Characterization of prevalence encoding

To generate a running estimate of the local sequence prevalence in the animal’s behavioral choices, we constructed, for each behavioral session, a vector of binary values of the same length as the session. Trials corresponding to the last step of the target sequence were given the value of ‘1’ (regardless of the context), all other trials were given the value of ‘0’. This binary vector was then convolved with a causal half-Gaussian kernel with zero mean and standard deviation of 20 trials.

We used five-fold cross-validated performance of a linear model to characterize the extent to which sequence prevalence factored into the activity of ACC neurons, using built-in Matlab functions ‘fitlme’ and ‘crossval’, and separately fitting the data in each behavioral context. Linear models were fit independently for each of the five analysis windows. For reporting fractions of cells with significant explained variance in each window, all five values for each neuron were used. For reporting explained variance of individual neurons, best of five values was used.

We defined explained variance as the coefficient of determination (R-squared) using the equation 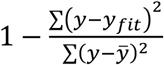 (equation 1), where the top is the sum of squared errors and the bottom is the total variance. The explained variance was calculated for ‘test’ subsets of data for each of the five cross-validation rounds, and then averaged across the five resulting values. In some cases, the explained variance was negative reflecting overfitting (typically occurred in the data sets with a small sample size). These negative values were replaced with zeros for subsequent reporting.

#### Interaction of prevalence and context

We considered several linear mixed-effect models to examine how a potential interaction of prevalence and context might account for the observed variance in sequence representation. The models considered incorporated sequence prevalence as a fixed effect predictor and could include a random intercept that varied by context. We used each of these features alone or together to fit the different models, and similarly estimated performance as the fraction of explained variance with fivefold stratified (with respect to context) cross-validation.

#### Comparison with models relating movement vigor and trajectory to neural activity

We resolved whether the observed modulation of ACC neural activity could be explained by variation in sequence timing or spatial trajectory by evaluating how well linear models using these parameters as regressors performed relative to the models above. For a measure of movement vigor, we calculated the time between first step ‘center in’ and last step ‘side in’; for the spatial trajectory, we first took snippets of the X and Y coordinates within the same 5 analysis windows that were used for all analyses of neural activity and then concatenated those snippets for all windows and both coordinates together in order to get a 1D vector. Then we combined such vectors for all the sequence instances within a session into a matrix and performed PCA to reduce the dimensionality of the raw spatial data but preserve the data about variation in animal’s motion. The first principal component was then used as a regressor for the corresponding linear model.

#### Decoding sequence prevalence from ACC ensemble trajectories

To characterize the extent, to which sequence prevalence could be decoded from population activity buy a downstream decoder, we evaluated the 5-fold cross-validated performance of several linear models that used the firing rates (or more specifically, the number of spikes) of individual neurons in each of the five analysis windows as regressors to estimate the local prevalence of the sequence in question during the t-th instance of sequence execution. The most general model (equation 2) modeled the response variable (local sequence prevalence at the time when the t-th sequence iteration was executed as a linear sum of predictor variable set, with each neuron contributing 5 spike counts, each with its own regression coefficient (weight).

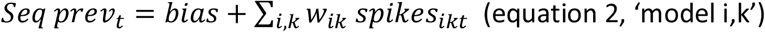

where t=sequence instance, i=1:number of neurons, k=1:5 analysis windows.

For this most general model, each neuron received a significance score for each of the five analysis windows that showed whether that particular neuron in that particular window contributes to the prediction of sequence prevalence. We used these scores to sort each neuron into classes 0 -5, depending on the total number of windows this neuron was significant in. Neurons that were not significant in any of the windows ended up in class 0 and were removed from all further modelling, and only sessions with at least 5 neurons outside of class 0 were retained for further analyses.

We further sometimes fit models on all neurons from class 1 through 5, and sometimes fit models only on neurons from classes 1 through 4. The latter was done to guarantee that we had removed those neurons that themselves had significant prevalence encoding in all windows, thus forcing the models to use multiple neurons to output good predictions the temporal extent of sequence execution.

To estimate the best possible prediction in each of the 5 analysis windows, we also fit 5 individual-window-models:

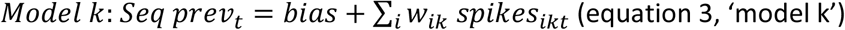

Finally, we fit a linear model that was constrained to always use the same readout (i.e. same weight for any given neuron in all five windows, equation 4, model *w*_*ik*_ = *w*_*il*_). In essence, this single aggregate model treats each of the five windows as extra datapoints for the same fit. As such, while having the same number of model parameters, this model benefits from these extra data but loses the temporal resolution, providing an estimate of how the fixed downstream decoder would perform on average during any instance of sequence execution.

